# Regulation of the transcriptome, miRNAs, and alternative splicing in a FSGS zebrafish injury model

**DOI:** 10.1101/2025.07.09.663814

**Authors:** Francescapaola Mattias, Olga Tsoy, Anna Iervolino, Stefan Simm, Elke Hammer, Alexander Gress, Tim Lange, Florian Siegerist, Sabine Ameling, Tim Kacprowski, Markus List, Olga Kalinina, Sören Franzenburg, Giovambattista Capasso, Karlhans Endlich, Jan Baumbach, Uwe Völker, Nicole Endlich, Felix Kliewe

## Abstract

**Background:** Focal Segmental Glomerulosclerosis (FSGS) is a severe kidney disorder with complex and not yet fully understood pathogenesis. Alternative splicing (AS) – the generation of distinct protein isoforms from the same gene – might play a critical role by the regulation of gene functions and disease development.

**Methods:** To investigate the role of AS in FSGS, we used a zebrafish model, which mimics key human FSGS features, including foot process effacement, matrix accumulation, podocyte detachment and parietal epithelial cell activation.

We performed total RNA sequencing of isolated zebrafish glomeruli and whole larvae, followed by integrative bioinformatic analysis to identify AS events and regulatory miRNAs.

**Results:** Our data revealed a downregulation of essential podocyte genes (*nphs1, nphs2, podxl, wt1*) and an inhibition of pathways associated with nephron development and cytoskeletal organization. We also observed increased expression of the transcription factor *stat3* and disease-associated miRNAs such as miR-21 and miR-193. AS analysis identified approximately ∼7,000 splicing events, primarily exon skipping (∼80%), affecting genes such as *nphs1*, *magi2*, and *ptpro*. A total of 136 and 612 alternatively spliced genes were found at 5 and 6 days post-fertilization (dpf), respectively. Isoform switch analysis uncovered 70 genes affected by AS in FSGS, including *epb41l5* (linked to podocyte adhesion), *fgfr1a* (fibroblast growth signaling), and members of the SRSF splicing factor family (e.g., *srsf3a*).

**Conclusions:** These findings emphasize the importance of transcriptional and post-transcriptional regulation, including AS, in FSGS pathogenesis. Furthermore, they support the zebrafish model as a valuable system for identifying novel mechanisms and potential therapeutic targets for kidney diseases.

## Introduction

Focal segmental glomerulosclerosis (FSGS) is a common histopathological pattern associated with nephrotic syndrome and progression to end-stage renal disease (ESRD) in both children and adults. Epidemiological studies have indicated that the global prevalence of FSGS has steadily increased over recent decades.^1–3^

Histologically, FSGS is characterized by the presence of scarring affecting a subset of glomeruli (focal), while others remain intact. Moreover, these scars are confined to specific segments of the glomerulus (segmental),^4,5^ leading to a reduction in the kidney’s filtration capacity and resulting in proteinuria. In severe cases, disease progression can culminate in renal failure.

The primary site of injury in FSGS is the podocyte, a post-mitotic epithelial cell that is crucial for maintaining the integrity of the glomerular filtration barrier.^6^ Studies, such as those conducted by Fornoni et al., have demonstrated that podocyte damage is associated with recurrent proteinuria in allograft kidneys of patients with idiopathic FSGS, as observed via electron microscopy.^7^

To date, kidney biopsy remains the gold standard for diagnosing FSGS.^8^ Due to its invasive nature, there is an urgent need for minimal-invasive diagnostic approaches based on novel biomarkers.

In this context, regulatory mechanisms like non-coding RNAs (ncRNAs) or alternative splicing (AS) has been identified as potential key factor in the pathogenesis of kidney disease and could lead to the definition of novel biomarker. Although research in this domain remains limited, recent investigations have explored the role of AS in different glomerular disease models and in podocytes subjected to mechanical stress.^9,10^

Mutations at splicing sites, alterations in splicing factors, and the creation or loss of splicing sites can disrupt pre-mRNA processing, leading to the production of non-functional or pathogenic proteins that contribute to disease manifestation.

To investigate the role of AS as well as ncRNAs, total RNA sequencing (RNA-Seq) is a powerful technique for investigating the transcriptomic landscape of scarred glomeruli, enabling the exploration of AS events, mutations, and RNA editing.

Due to ethical and practical constraints, zebrafish larvae (*Danio rerio*) are a widely accepted model organism in kidney research. Their pronephros which consists of a single glomerulus and two kidney tubules exhibits high structural and transcriptomic homology with the human kidney. Previously, we established an FSGS model in zebrafish larvae.^11–13^ This model exhibited podocyte depletion, activation of parietal epithelial cells (PECs), and thickening of the glomerular basement membrane (GBM).^11^ These findings are consistent with studies by Lee, Kuppe, and Dijkman.^14–16^ Furthermore, zebrafish larvae have been used to develop high-content drug screening assays, resulting in the identification of Belinostat as a protective agent in the FSGS zebrafish model.^12^

In this study, Nifurpirinol (NFP) was used to induce podocyte-specific ablation, thereby simulating FSGS in zebrafish larvae. Subsequently, we analyzed the full transcriptome in whole zebrafish larvae and of isolated zebrafish glomeruli using different RNA-Seq techniques to combine the epigenetic regulatory factors, focusing on miRNAs and AS events, to identify interaction partners and potential regulatory networks. Our study provides novel insights into FSGS pathogenesis and paves the way for the identification of novel disease biomarkers and targeted therapeutic strategies.

## Methods

### Zebrafish strains and husbandry

Zebrafish were handled and maintained as previously described.^17^ The stages are indicated in days post-fertilization (dpf). For this work, the zebrafish Tg(nphs2: GAL4-VP16); Tg(UAS:Eco.nfsB-mCherry) strain was used. Larvae express the bacterial enzyme nitroreductase and the fluorescent protein mCherry exclusively in podocytes. All experiments were conducted in accordance with the German animal protection law, were overseen and approved by the “Landesamt für Landwirtschaft, Lebensmittelsicherheit und Fischerei, Rostock” (LALLF M-V) of the federal state of Mecklenburg-Western Pomerania.

### Experimental design of the FSGS zebrafish injury model

Larvae at 4 dpf were first selected for a strong homogeneous mCherry using a fluorescence stereomicroscope (SMZ 18 Nikon, Düsseldorf, Germany). FSGS was induced by the treatment of the zebrafish larvae with 50 nM Nifurpirinol (Sigma Aldrich, St. Louis, MO, USA) dissolved in 0.1% dimethyl sulfoxide (DMSO, Sigma Aldrich) in E3 medium. 0.1% DMSO was used for control vehicles as described previously.^11^ A treatment period of 24 hours was maintained for all experiments. Larvae and isolated glomeruli were collected at 24-and 48-hours post-treatment, corresponding to 5 and 6 dpf, respectively. Treatment with 50 nM Nifurpirinol resulted in the development of periocular and abdominal edema in the transgenic larvae. Furthermore, treatment induced podocyte damage, as indicated by a reduction in mCherry fluorescence and its subsequent accumulation in the proximal tubule (Fig. S1).

### Larvae collection and glomeruli isolation

NFP- and DMSO-treated larvae at 5 and 6 dpf were dissolved in 1 ml Tri-Reagent (Sigma-Aldrich) according to manufacturer’s instructions and homogenized with ceramic beads (Carl Roth, Karlsruhe, Germany) at 5 m/s for 20 s. Glomeruli of NFP-treated and untreated larvae were isolated manually at 5 and 6 dpf using two extra-fine precision tweezers (DUMONT, Montignez, Switzerland) and a fluorescence stereomicroscope (SMZ 18 Nikon, Tokio, Japan). Pools of 10 glomeruli were collected and centrifuged at 1000 x g for 5 min at 4°C. Subsequently, the supernatant was discarded and 50 µl of Tri-Reagent (Sigma-Aldrich) were added. Whole larvae and isolated glomeruli were used for the total RNA-Seq and AS analyses.

### RNA extraction, cDNA synthesis and qRT-PCR

For total RNA, samples were processed in Tri-Reagent (Sigma-Aldrich) according to manufacturer’s instructions. For cDNA synthesis, 1 µg of the isolated total RNA was transcribed using the QuantiTect Reverse Transcription Kit (Qiagen, Hilden, Germany). The quantitative real-time PCR (qRT-PCR) analysis was performed on a Bio-Rad iQ5 (Bio-Rad, Hercules, CA, USA) using the iTaq Universal SYBR Green Supermix (Bio-Rad) with *ef1a* as reference gene. Relative quantification of the mRNA levels was done by the efficiency corrected calculation model by Pfaffl^18^ and values are shown with standard deviations (SD) or standard error of the mean (SEM) from at least three biological replicates. Used primers are summarized in Supplemental Table 1.

### miRNA isolation and sequencing

MicroRNAs were isolated using the miRNeasy Micro Kit (Qiagen), which is specifically optimized for low-input samples. RNA concentration and integrity were evaluated using the Agilent Bioanalyzer system with the RNA 6000 Pico Kit (Agilent Technologies, Waldbronn, Germany). Sequencing of miRNAs was performed by GenXPro (Frankfurt am Main, Germany) and combined with the mRNA technique MACE (Massive Analysis of cDNA Ends) methodology, which enables digital expression profiling with high sensitivity and accuracy.

### RNA-Seq and transcriptome & alternative splicing analysis

Samples were processed in Tri-Reagent (Sigma-Aldrich) according to the manufacturer’s instructions, followed by DNA-digestion steps using the RNase-Free DNase I Kit and RNA Clean-Up and Concentration kit according to manufacturer’s instructions (both Norgen Biotek Corp., Thorold, ON, Canada). Briefly, after quality assessment using a Tape Station 4200 (Agilent, Santa Clara, CA, USA), a PolyA library was prepared based on 500 ng total RNA per sample using the TruSeq stranded mRNA kit (Illumina, San Diego, CA, USA). Subsequently, for sequencing 200 pM of the final library were loaded on a NovaSeq 6000 System, S1 FlowCell (2 x 100 bp pair-end reads). A sequencing depth of ∼ 200 Mio clusters per sample was obtained and recorded at the Competence Centre for Genomic Analysis in Kiel. RNA-Seq raw reads for NFP-treated zebrafish larvae and isolated glomeruli after both 5 and 6 dpf and controls were mapped to the zebrafish reference genome GRCz11 (Genome Reference Consortium Zebrafish Build 11) with the genome annotation from Ensembl version 104 using STAR version 2.7.5c.^19^ We used the default parameters with a 2-pass mode recommended for AS analysis. Gene read counts were obtained by the featureCounts tool included in Subread 2.0.0.^20^ Normalization and differential expression analysis were carried out using DESeq2 version 1.38.3.^21^ We defined significantly differentially expressed genes with p-value < 0.05 after the Benjamini-Hochberg correction. We visualized the differentially expressed genes using the EnhancedVolcano R package.^22^ The heatmaps of differentially expressed genes were visualized with seaborn.clustermap function. They include on the Y-axis a dendrogram of the hierarchical clustering with a Euclidian distance of normalized and z-score standardazed gene expression values in the different samples. The gene enrichment analysis was performed using Gene Set Enrichment Analysis (GSEA) with Gene Ontology terms.^23^ The dotplots of the overrepresented categories from the GO biological processes were created using the library clusterProfiler.^24^

For isoform switch analysis, we prepared transcript counts using salmon version 1.9.0^25^ with the transcript annotation based on Ensembl version 104 and ran the R package IsoformSwitchAnalyzeR^26^ version 1.20.0 with the DESeq2 algorithm. For alternative splicing analysis, we used leafcutter version 0.2.9,^27^ rMATS version 4.1.2,^28^ and Whippet version 0.11.1 (with julia 1.6.6)^29^ with the default parameters. We defined significantly differentially used events as events with a p-value < 0.05 (for leafcutter and rMATS) or a confidence > 0.95 (for Whippet).

### Laser scanning microscopy

Images were captured with an Olympus FV3000 confocal microscope (Olympus, Tokyo, Japan) with 20x/40x/60x oil immersion objectives and Olympus FV3000 CellSense software.

### Single-cell RNA-sequencing data processing and analysis

Gene expression matrices derived from glomerular single-cell RNA-sequencing experiments (control, nephrotoxic serum nephritis (NTS) day 1 and day 5, and doxorubicin nephropathy) were downloaded from the BroadSingleCell Portal^30^ and imported into R (v4.2.0) and processed using the Seurat package (v5.0.1). Each sample was converted into a Seurat object using the CreateSeuratObject function with a minimum of 200 genes and 3 cells per feature. Cells were annotated by sample origin and merged to generate combined datasets for NTS (control, NTS day 1, and NTS day 5) and doxorubicin (control and doxorubicin) analyses. Quality control included filtering cells with <200 or >5,000 detected genes, <1,000 or >10,000 UMIs, or >5% mitochondrial content. Following normalization (NormalizeData), variable features were identified, and datasets were scaled with the ScaleData function. Principal component analysis (PCA) was conducted on variable genes, followed by dimensionality reduction via UMAP (RunUMAP) and clustering (FindNeighbors, FindClusters). Clusters were manually annotated using canonical glomerular cell markers (e.g., *Nphs1* (podocytes), *Pecam1* (endothelial cells), *Pdgfrb* (mesangial cells), *Adgre1* (leukocytes), *Cldn1* (parietal epithelial cells)). Subclusters of podocytes (POD), mesangial cells (MC), and endothelial cells (EC) were extracted and reanalyzed using the same clustering approach. Differential gene expression between clusters was determined using the FindAllMarkers function with thresholds of log₂ fold change > 0.25 and minimum expression in 25% of cells. Data visualization included UMAPs, violin plots, feature plots, and heatmaps which were all generated in Seurat. Marker genes and expression profiles were exported as CSV files for further downstream analysis described under “statistical analysis”.

### Statistical analysis

The GraphPad Prism 9 software was used for statistical analysis of experimental data and preparation of graphs. Scatter plots indicate individual units used for statistical testing (samples or replicates), as specified in respective figure legends. Data are presented as mean ± SD (or SEM, as indicated). Statistical analysis was performed using an unpaired two-tailed t-test (n = number of independent experiments). For multiple groups statistical analyses were done by ANOVA with a Benjamini-Hochberg post-hoc test. Statistical significance was defined as p < 0.05 and significance levels are indicated as * p < 0.05, ** p < 0.01, *** p < 0.001, **** p < 0.0001 and non-significant (ns) in respective figure panels. The number of independent experiments and analyzed units are stated in the figure legends.

## Results

### Transcriptome analysis of zebrafish larvae and isolated zebrafish glomeruli after FSGS-induction

The present study employed the well-established Nifurpirinol-induced FSGS zebrafish larvae model (Fig. S1).^11^ RNA-Seq analysis revealed significant alterations in gene expression following NFP treatment in zebrafish larvae at 5 and 6 dpf.

At 5 dpf, 230 genes were identified as differentially expressed (DEGs), including 12 downregulated and 218 upregulated transcripts showing at least a two-fold change in expression. This number increased markedly at 6 dpf, with 770 DEGs detected, comprising 231 downregulated and 476 upregulated genes (Fig. 1A-B). This substantial increase in DEGs at 6 dpf indicates a progressive transcriptional response to NFP exposure (Fig. 1A-C & Fig. S2). While 24% of all DEGs were common to both time points, 28% were uniquely expressed at 5 dpf and 48% were specific to 6 dpf, suggesting a time-dependent regulation of gene expression (Fig. 1C). Hierarchical clustering was performed to assess sample correlation, revealing that gene expression profiles are strongly associated with NFP-treatment (Fig. 1D-E). Figure 1F shows the top 10 up- and down-regulated genes (sorted and selected by log_2_ fc). In the gene set enrichment analysis of Gene Ontology (GO) terms for biological process (BP), molecular function (MF) and cellular compartment (CC) we observed differences between the up- and down-regulated DEGs within the 5 dpf and 6 dpf as well as between them (Fig. S2). After NFP-treatment, down-regulated DEGs showed an enrichment in Gene Ontology (GO) terms for “gene expression regulation” and “chromatin dynamics” at 5 dpf, while up-regulated DEGs were enriched in “hormonal and cytokine signaling pathways” (Fig. 1G). In contrast, 48 hours post-treatment (6 dpf), down-regulated DEGs were enriched in “immune response and homeostasis maintenance”, and up-regulated DEGs enriched in “metabolic pathways” (Fig. 1H).

**Fig. 1:**
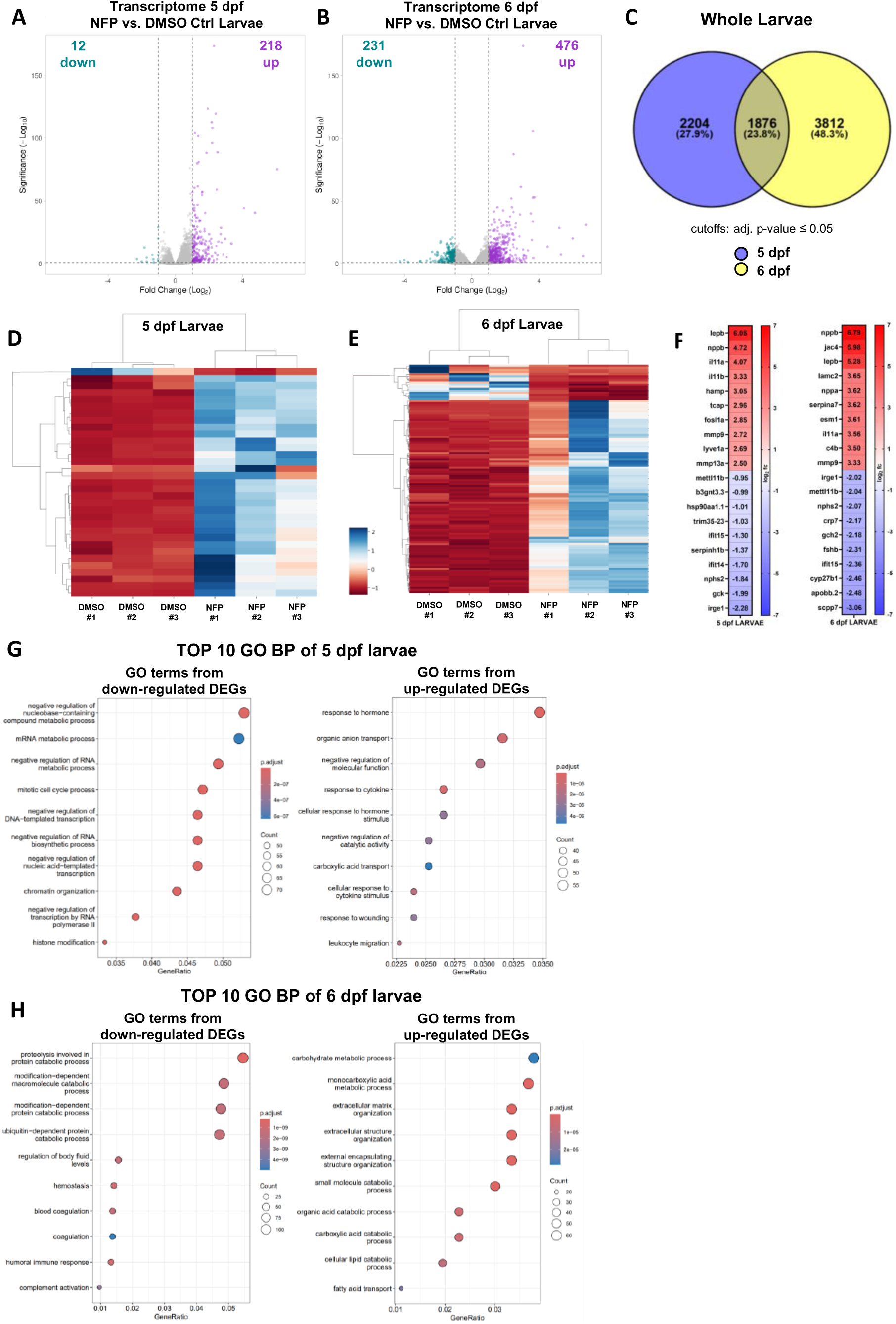
Transcriptomic analysis of zebrafish larvae. (A, B) Volcano plot of differentially expressed genes (DEGs) in NFP-treated zebrafish larvae at 5 dpf (A) and 6 dpf (B) compared to the control group. The purple dots and the cyan dots represent significant up-regulated and down-regulated DEGs (log2 fold-change ≥ 1; adjusted p-value ≤ 0.05). The grey dots represent genes with no significant differences. (C) Venn diagram shows the significantly DEGs at 5 and 6 dpf after NFP treatment and their overlap (adj. p-value ≤ 0.05). (D, E) Heatmaps of gene expression between NFP- and DMSO-treated larvae at 5 dpf (D) and 6 dpf (E) based on hierarchical clustering analysis. Heatmaps colors represent relative gene expression as indicated in the color key. (F) Heatmaps show the top 10 up- and down-regulated genes present in zebrafish larvae at 5 and 6 dpf that show significant changes in gene level due to NFP treatment. Data are expressed as log2 fold change, a p-value < 0.05 was considered as significant. Red: up-regulated; blue: down-regulated compared to controls. (G, H) The top 10 biological processes (BP) enriched in DEGs of NFP- treated larvae at 5 (G) and 6 (H) dpf compared to the respective control. The size of the dots represents the number of genes and the color of the dots represent the adjusted p-values.

Additionally, we performed a similar analysis for glomeruli isolated from the NFP-treated zebrafish larvae at 5 and 6 dpf as well as the DMSO-treated controls. Volcano plots were used to visualize DEGs (adj. p-value ≤ 0.05 and a minimum two-fold change in expression) (Fig. 2A-B). At 5 dpf, 1,007 DEGs were identified (578 down-, 429 upregulated), and at 6 dpf, 2,299 DEGs (1,042 down-, 1,257 upregulated) (Fig. 2A-B). An overlap of 22% of all DEGs was observed between the time points, while 15% were unique to 5 dpf and 63% unique to 6 dpf (Fig. 2C). Hierarchical clustering revealed distinct DEG patterns between NFP-treated and control groups at both time points (Fig. 2D-E). Within the top 10 up- and down-regulated genes we found *nphs1*, *nphs2*, *ptpro*, *lepb* and *ntng2b* (Fig. 2F). After NFP-treatment, downregulated DEGs showed an enrichment in GO BP terms related to "nephron and glomerulus development", while up-regulated DEGs were linked to "RNA and ribosome activities" and “mitochondrial gene expression” (Fig. 2G, Fig. S3 and Fig. S4).

**Fig. 2:**
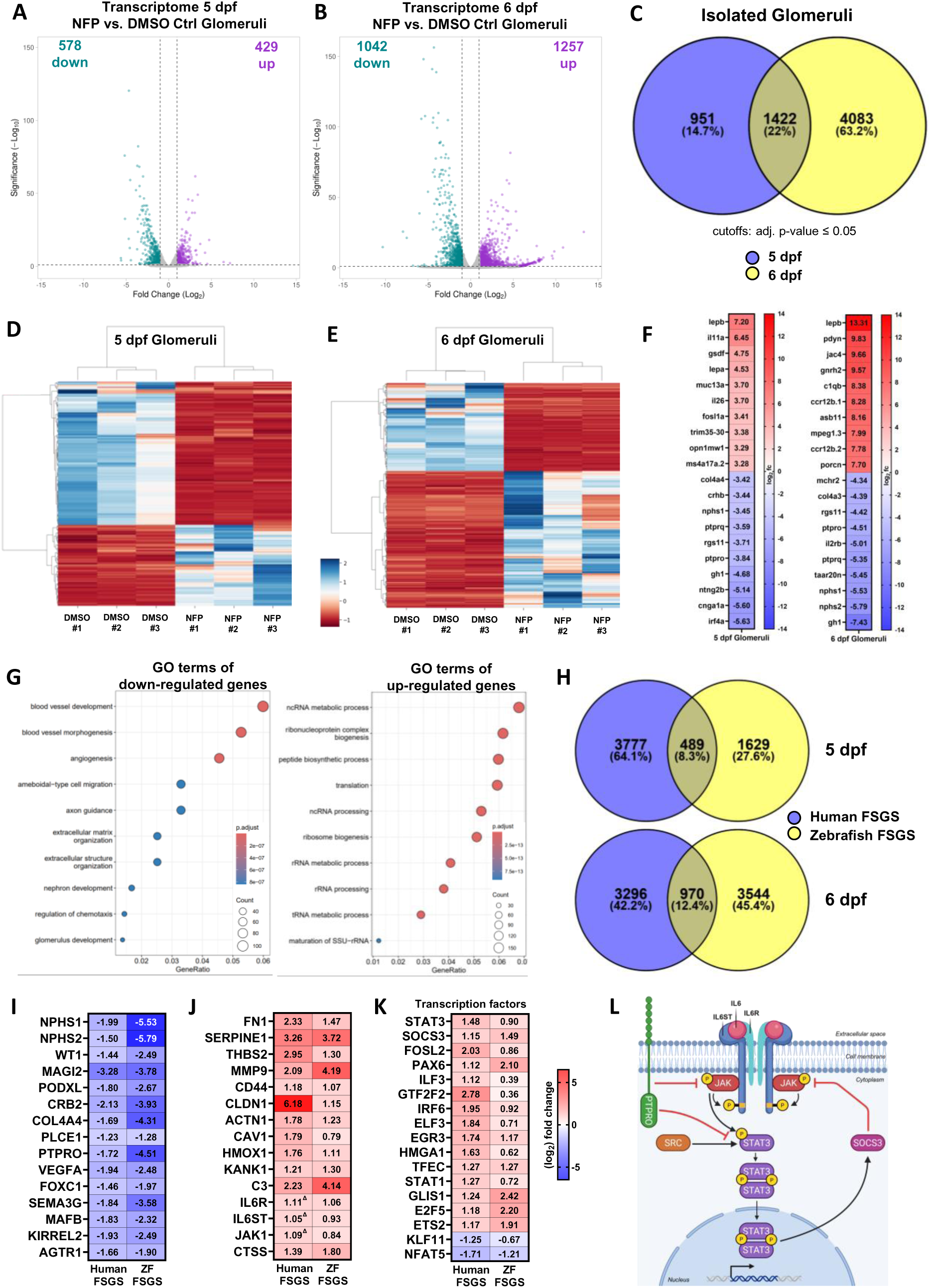
**Transcriptomic analysis of isolated zebrafish glomeruli.** (A, B) Volcano plot showing differentially expressed genes (DEGs) in isolated glomeruli of zebrafish larvae at 5 dpf (A) and 6 dpf (B) (NFP vs. DMSO control). The purple dots and the cyan dots represent significantly up- and down-regulated DEGs (log2 fold-change ≥ 1; adjusted p-value ≤ 0.05). The grey dots represent genes with no significant differences. (C) Venn diagram depicts the number of the common and distinct DEGs at 5 and 6 dpf in isolated glomeruli (NFP vs Ctrl) (adj. p-value ≤ 0.05). (D, E) Heatmaps of hierarchical clustering for gene expression in isolated glomeruli at 5 dpf (D) and 6 dpf. Heatmaps colors represent relative gene expression as indicated in the color key. (F) Heatmaps of the top 10 most significantly up- and down-regulated DEGs at 5 and 6 dpf. Data are expressed as log2 fold change, a p-value < 0.05 was considered as significant. Red: up-regulated; blue: down-regulated compared to controls. (G) Top 10 enriched biological processes for DEGs at 6 dpf based on Gene Ontology analysis. Dot size represents gene count; color indicates p-adjusted values. (H) Venn diagrams showing shared DEGs between human FSGS glomeruli and zebrafish glomeruli at 5 dpf and 6 dpf. Human FSGS dataset from www.nephroseq.de “Hodgin FSGS Glom Dataset”.^31^ (I-K) Heatmaps depicts significantly downregulated (I) and upregulated (J) genes, as well as putative FSGS-associated transcription factors (K) identified in both human FSGS patients and in our zebrafish model of FSGS at 6 days post fertilization (dpf). Data shows glomerular expression as log2 fold change (ZF FSGS) or unlogged fold change (human FSGS) (Δ: non-significant). (L) Schematic representation of the IL6/STAT3 signaling pathway under FSGS conditions. Upregulation of IL6R, IL6ST, and JAK1 leads to enhanced STAT3 activation via phosphorylation. The phosphorylated form (pSTAT3) translocates to the nucleus and induces expression of downstream targets such as SOCS3. The downregulation of PTPRO impairs dephosphorylation of STAT3.

### Cross-species analysis of FSGS-related genes in zebrafish and human

To further validate our model, we compared RNA-Seq data from zebrafish glomeruli with microarray data from FSGS patient glomeruli.^31^ At 5 dpf, 489 DEGs (8%), and at 6 dpf, 970 DEGs (12%), were shared between the two datasets (Fig. 2H & Supplemental Data 2). Our FSGS zebrafish injury model revealed significant alterations in a large number of known genes, which are regulated in human FSGS patients (Fig. 2I-J). Detailed analysis of downregulated genes showed association with kidney, glomerulus, and nephron development, highlighting key genes such as *nphs1*, *nphs2*, *wt1*, *magi2*, *podxl*, *crb2*, *plce1,* and *ptpro* (Fig. 2I). Key podocyte-specific transcription factors such as *foxc1* and *mafb* were also significantly downregulated in FSGS-induced zebrafish glomeruli (Fig. 2I). In contrast, the expression of other potential FSGS-related transcription factors, such as STAT3 – whose activation in podocytes is critical for the development of glomerulonephritis^32^ – was significantly increased (Fig. 2K), thereby promoting the upregulation of SOCS3 (Fig. 2K). Single-cell RNA sequencing (scRNA-Seq) published by Chung et al.^30^ revealed a significant upregulation of *Stat3* and *Socs3* expression in podocytes in both nephrotoxic serum nephritis (NTS) or doxorubicin (DOX)-induced models of kidney injury (Fig. S5B-C). Other genes relevant for podocytes function and stability are significantly upregulated in FSGS patients as well as in the FSGS zebrafish model, such as *FN1*, *SERPINE1*, *THBS2*, *ACTN1*, and *KANK1* (Fig. 2J). Furthermore, *IL6R* and *IL1B*, which play a pivotal role in inflammatory processes, exhibited a significant increase in expression in FSGS-induced zebrafish glomeruli (Fig. 2J). Upregulation of *IL6R*, *IL6ST*, and *JAK1* suggests enhanced *STAT3* activation via phosphorylation and nuclear translocation, promoting expression of targets such as *SOCS3* (Fig. 2L). Concurrently, the downregulation of *PTPRO* in FSGS-induced glomeruli (Fig. 2I) as well as in NTS- and DOX-induced models of kidney injury (Fig. S5A) impairs the dephosphorylation and inactivation of STAT3, thereby sustaining its activity (Fig. 2L).

### Regulation of miRNA in the FSGS zebrafish injury model

In our total RNA-Seq analysis, we identified genes with known connections to miRNA regulation. This prompted us to examine miRNA expression in more detail. To this end, we analyzed the expression profiles of miRNAs in isolated glomeruli from zebrafish subjected to FSGS-induced injury.

Differential expression analysis of miRNAs in isolated glomeruli from the FSGS zebrafish model compared to controls revealed a total of 42 differentially expressed miRNAs (Fig. 3A-B). In addition to the well-characterized miR-21 and miR-193, we identified novel potentially relevant miRNAs, including miR-146, miR-34, miR-200, and miR-429, all of which exhibited significantly increased expression levels (Fig. 3A-B).

**Fig. 3:**
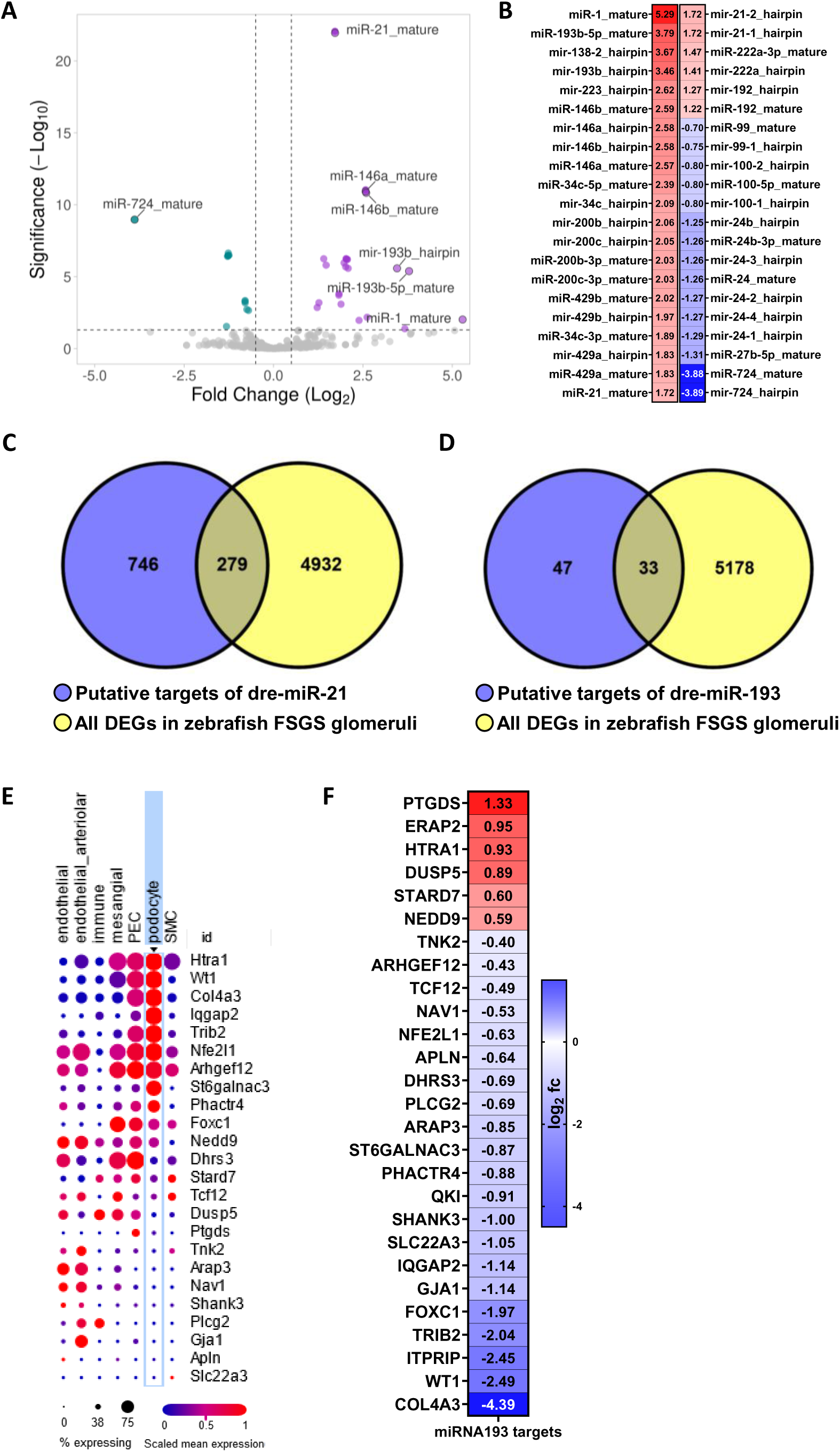
**Differential expression of miRNAs in glomeruli of FSGS-induced zebrafish larvae.** (A) Volcano plot of differential expression of miRNAs in isolated glomeruli after podocyte-specific glomerular injury compared to control groups. Data represent miRNA expression profiles identified by RNA sequencing. Significantly upregulated (purple) and downregulated (cyan) miRNAs are highlighted (adjusted p-value < 0.05; log2 fold change ≥ 0.5). (B) Heatmap of all significantly up- and down-regulated miRNAs after FSGS induction. (C) Overlap of putative targets of miR-21 (taken from miRDB^75^) with all DEGs in zebrafish glomeruli after FSGS induction. (D) Overlap of putative targets of the dre-miR-193 (taken from Gebeshuber et al.^34^) with all DEGs in zebrafish FSGS glomeruli. (E) Single-cell RNA-seq data of mouse glomerulus (published by Chung et al.^30^) from all putative miR-193 targets found in the zebrafish injury model. (F) Heatmap of the glomerular expression patterns of miR193 targets in NFP-treated zebrafish larvae. Data are expressed as log2 fold change, a p-value < 0.05 was considered as significant. Red: up-regulated; blue: down-regulated compared to controls.

Due to their well-established roles in chronic kidney disease (CKD) and focal segmental glomerulosclerosis (FSGS), subsequent analyses were focused on the upregulated miR-21 and miR-193.^33,34^ mRNA-target prediction of miR-21 with all mRNA DEGs in zebrafish glomeruli after FSGS induction generated 279 potential targets (Fig. 3C) including *epb41l5, npr3* or *ptpro* (Supplemental Data 2). For miR-193, 33 potential targets were differentially expressed in zebrafish glomeruli after FSGS induction (Fig. 3D-E and Supplemental Data 2). Using published single cell RNA-Seq from mouse^30^ allowed us to check the expression level of the 33 potential miR-193 targets within the different cell types of kidney. *Htra1*, *Wt1*, *Col4a3*, *Iqgap2*, *Trib2*, *Nfe2l1*, and *Arhgef12* showed a high expression in podocytes (Fig. 3E). Supporting this, key targets of miR-193 implicated in glomerular diseases, such as *wt1*, *col4a3*, and *foxc1*, were significantly downregulated in glomeruli isolated from NFP-treated larvae (Fig. 3F).

### Alternative splicing events in FSGS-induced zebrafish larvae

Besides the regulation due to miRNAs a second important regulatory mechanism on the molecular level is the generation of AS isoforms. For the identification of alternatively spliced genes, splicing events were detected using rMATS, LeafCutter, and Whippet. Additionally, isoform switches were analyzed with IsoformSwitchAnalyzeR, and all resulting genes were subsequently screened for podocyte-specific expression (Fig. 4A). We used sankey analysis to visualize the distribution of splicing events (Fig. 4B). A comprehensive list of all identified AS genes is provided in Supplementary Data 3. Quantification of the frequency of splicing event classes (including exon skipping, intron retention, and alternative splice sites) in isolated glomeruli and whole zebrafish larvae showed that exon skipping comprise 79-86% (Fig. 4C). Combining the identified AS events from all tools and assigning them to the zebrafish gene annotation lead to the identification of 97 (larvae) and 136 (glomeruli) genes with potential AS isoforms in zebrafish at 5 dpf (Fig. 4D, F and Supplemental Data 2). In contrast, at 6 dpf, the number of potential AS isoform genes increased by a factor of ∼3 (242 AS genes) in whole larvae and a factor of ∼5 (612 AS genes) in glomeruli (Fig. 4E, G). The overlap of the genes between both time points was 21 (larvae) and 47 genes (glomeruli), suggesting that a subset of splicing events remains consistent across developmental stages (Fig. S6A-B; Fig. S7; Supplemental Data 2).

**Fig. 4:**
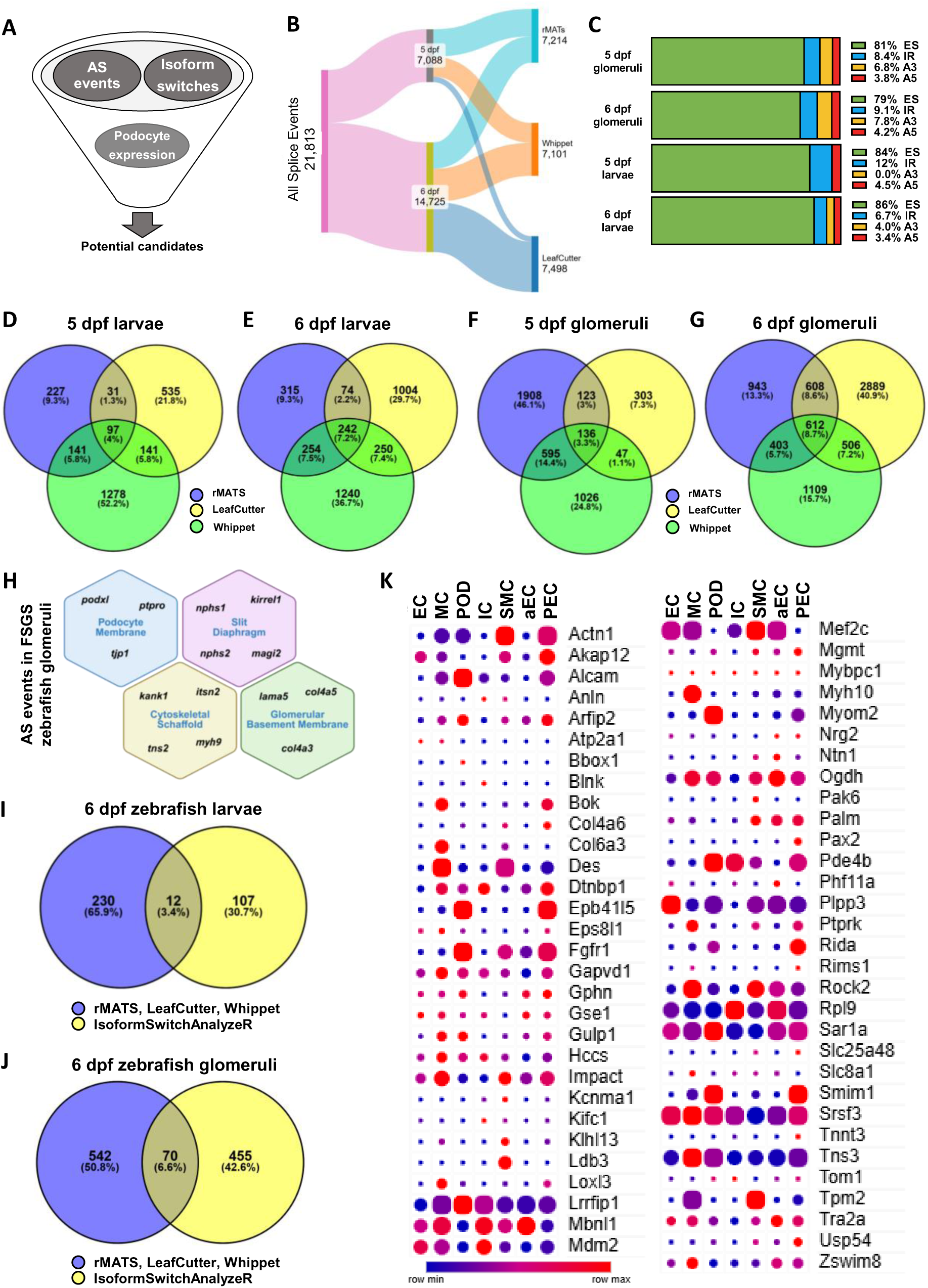
**Differential alternative splicing events in glomeruli of zebrafish larvae.** (A) Methodical workflow for the identification of potential candidate genes. Alternative splicing (AS) events were identified using rMATS, LeafCutter, and Whippet. Additionally, isoform switches were analyzed with IsoformSwitchAnalyzeR. Finally, all identified genes were screened for podocyte-specific expression. (B) Sankey’s diagram illustrates the flow of alternative splicing events through three tools (rMATS, LeafCutter, Whippet) for isolated 5 dpf and 6 dpf glomeruli. (C) Percentage distribution of the types of AS events found in rMATS: ES (exon skipping); IR (intron retention); A5SS (Alternative 5′ splice site); A3SS (Alternative 3′ splice site). (D-G) Venn diagram depicts the overlap of genes that showed a significant splice event at 5 dpf (24h after treatment) and 6 dpf (after 48h treatment) in rMATS, leafcutter and Whippet in larvae and isolated glomeruli. (H) Shown are primarily podocyte-specific genes with AS events detected in glomeruli upon FSGS induction, which are also known to play a role in glomerular disease. (I) The overlap shown in Fig. 4E was integrated with isoform switch analysis of whole larvae at 6 days post fertilization (dpf). (J) The intersecting genes from Fig. 4G were cross-referenced with isoform switching data from isolated glomeruli at 6 dpf. (K) Single-cell RNA-Seq expression data of mouse glomerulus (published from Chung et al.^30^) of all alternatively spliced genes found parallel in rMATS, LeafCutter, Whippet and IsoformSwitchAnalyzeR. EC, glomerular endothelial cells; MC, mesangial cells; POD, podocytes; IC, immune cells; SMC, smooth muscle cells; aEC, arteriolar endothelial cells; PEC, parietal epithelial cells.

We identified glomerular-specific alternatively spliced genes with established causal roles in glomerular disease, most of which are predominantly expressed in podocytes. These include genes encoding proteins localized to the podocyte membrane (e.g., *podxl*, *ptpro, tjp1*), the cytoskeletal scaffold (e.g. *tns2*, *myh9*, *itsn2*, *kank1*), the glomerular basement membrane (e.g. *lama5*, *col4a3*, *col4a5*), and the slit diaphragm (e.g. *nphs1*, *nphs2*, *kirrel1 or magi2*) (Fig. 4H and Supplemental Data 2).

To refine target selection, the next step was to identify alternatively spliced genes that also exhibited an isoform switch following FSGS induction. The IsoformSwitchAnalyzeR found 12 genes in larvae and 70 genes in glomeruli, which showed a significant isoform switch after FSGS induction (Fig. 4I, J). Next, we analyzed the glomerular expression pattern of these genes focusing on their expression in podocytes (Fig. 4K and Fig. S8). Notably, three candidate genes – *epb41l5*, *fgfr1a*, and *srsf3a* – met all criteria, including consistent AS detection, differential isoform usage, and podocyte-specific expression, underscoring their potential relevance in FSGS.

### Regulatory modes of alternative splicing isoform genes, including *epb41l5*, *fgfr1a*, and *srsf3a*, in glomeruli from FSGS-injured zebrafish larvae

Erythrocyte Membrane Protein Band 4.1 Like 5 (*epb41l5*), a key regulator of cell polarity and adhesion, is highly expressed in zebrafish and human glomeruli (Fig. S9A-B). However, *epb41l5* expression was significantly reduced after induction of FSGS at 5 and 6 dpf, consistent with findings in patients with FSGS^35^ (Fig. 5A and Fig. S9C). Four differentially abundant splice variants of *epb41l5* were identified in NFP-treated glomeruli compared to controls (Fig. 5B-C and Fig. S9D) and two of these with a specific isoform switch in FSGS-injured zebrafish glomeruli at 5 and 6 dpf (Fig. 5D and Fig. S9E). Both isoforms contain the FERM domain; however, isoform epb41l5-204 (ENSDART00000135334) lacked the C-terminal region, suggesting potential functional divergence from the other isoforms. Among them, epb41l5-202 (ENSDART00000101276) exhibited the highest expression and was significantly downregulated upon NFP treatment (Fig. 5C and Fig. S9D). Quantitative real-time PCR confirmed the differential expression of *epb41l5* isoforms in NFP-treated versus control glomeruli (Fig. 5E). Interestingly, we found that the *Epb41l5* interacting partners *Crb2* and *Mpp5* (*Pals1*) were also significantly downregulated after the induction of FSGS (Fig. 5F-G). Furthermore, scRNA-Seq^30^ also revealed a significant podocyte-specific downregulation of *Epb41l5, Crb2* and *Mpp5* in NTS- and DOX-induced models of kidney injury (Fig. 5H-J).

**Fig. 5:**
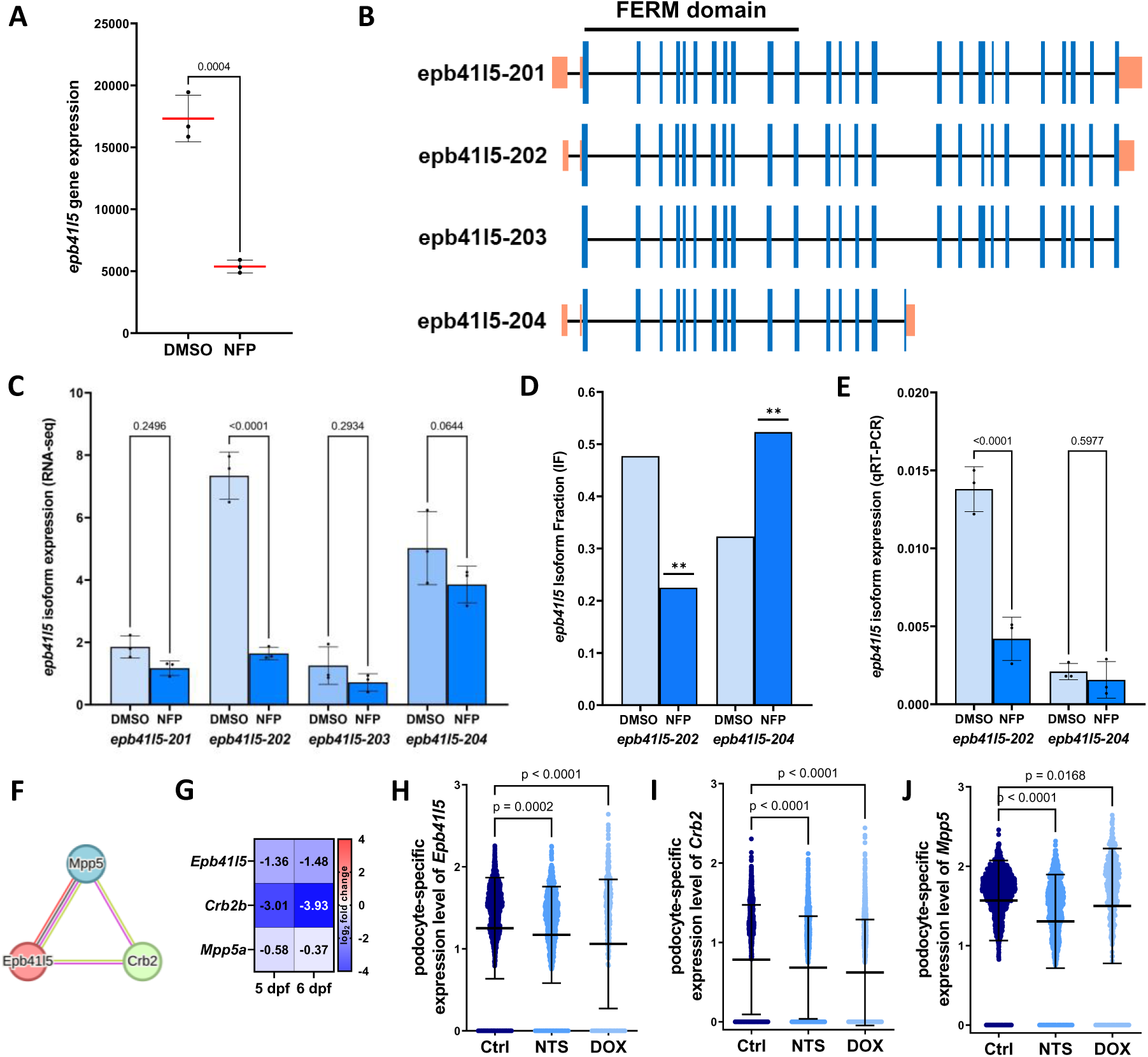
Regulation of *epb41l5* in FSGS-injured zebrafish glomeruli. (A) Gene expression of *epb41l5* in glomeruli treated with DMSO and NFP at 6 dpf. (B) Overview of all *epb41l5* isoforms found in zebrafish glomeruli. FERM domain is highlighted. Blue: coding exons; orange: untranslated regions (UTRs). (C) Expression of *epb41l5* isoforms based on DESeq2 normalized gene counts from RNA-Seq at 6 dpf. (D) *Epb41l5* isoform fraction based on IsoformSwitchAnalyzer data at 6 dpf. (E) Validation of *epb41l5* isoform expression by qRT-PCR normalized to Ctrl glomeruli (DMSO-treated) and *Gapdh*. (F) STRING protein–protein interaction network showing strong correlated interactions of epb41l5 with Crb2 and Mpp5. (G) Heatmaps of gene expression levels of *epb41l5* and his interaction partner CRUMBS2 (*crb2a*) and PALS1 (*mpp5a*). Data are represented as log2 fold change, a p-value < 0.05 was considered as significant. Red: up-regulated; blue: down-regulated compared to controls. (H-J) Single-cell RNA-Seq data shown podocyte-specific gene expression of *Epb41l5* (H), *Crb2* (I) and *Mpp5* (J) in mouse with nephrotoxic serum nephritis (NTS) at day 5 and doxorubicin (DOX) compared to the control (Ctrl) (Chung et al.^30^). All gene annotations used in this analysis were based on Ensembl release 113 (October 2024).

In contrast, the total mRNA expression of *fgfr1a* (fibroblast growth factor receptor 1a), a tyrosine kinase receptor, was not altered significantly in zebrafish glomeruli after FSGS induction (Fig. 6A-B), but analysis of *fgfr1a* transcripts revealed that two isoforms exhibited expression changes in zebrafish glomeruli (Fig. 6C-D). Fgfr1a-204 (ENSDART00000147742) was downregulated, whereas fgfr1a-201 (ENSDART00000024882) was upregulated (Fig. 6C-D). Protein structure analysis revealed that we have a homologous substitution of the domain that mitigates the interaction to the fibroblast growth factor (Fig. 6E-G).

**Fig. 6:**
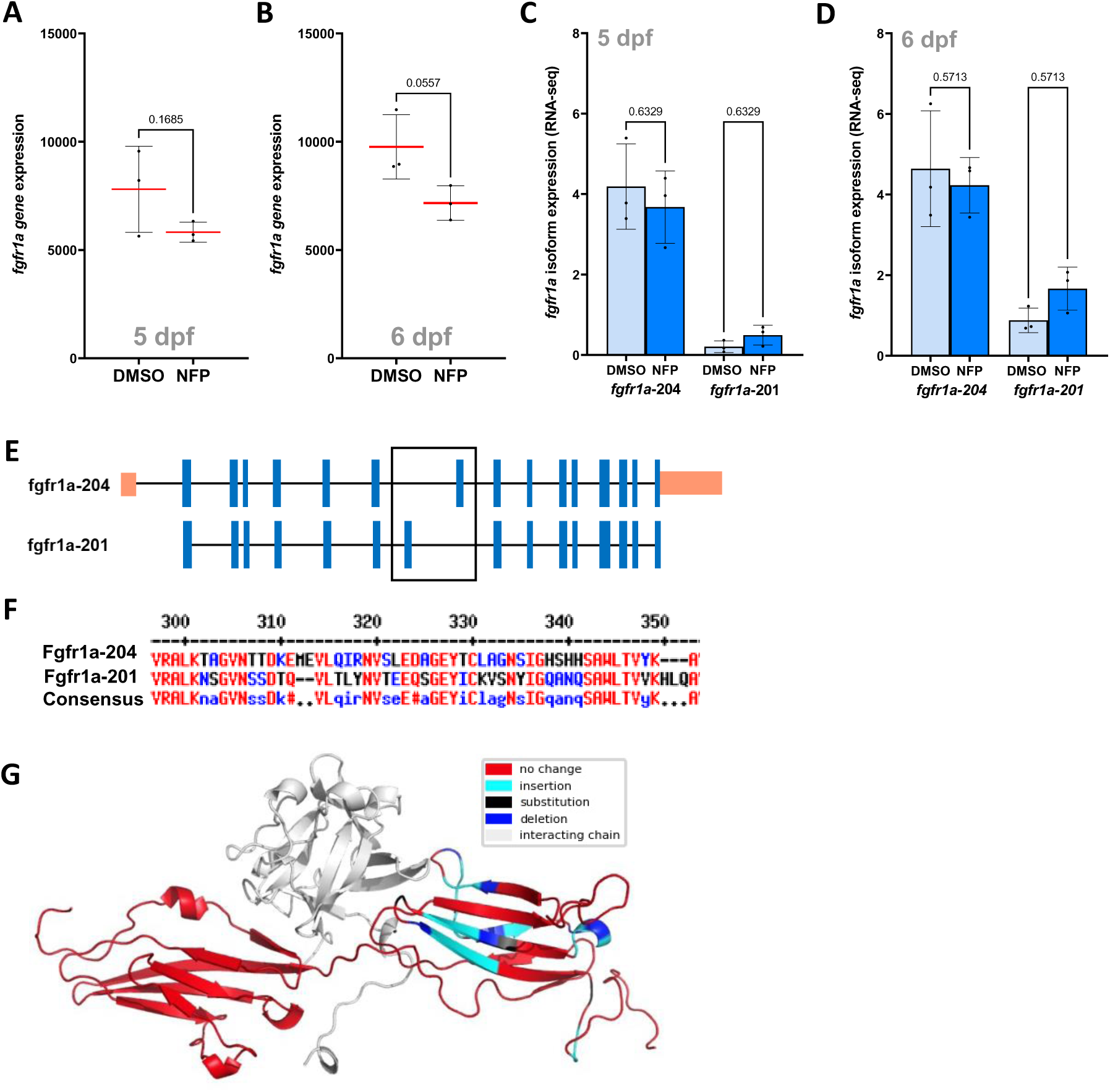
Regulation of *fgfr1a* in FSGS-injured zebrafish glomeruli. (A-B) *Fgfr1a* gene expression in DMSO and NFP glomeruli at 5 dpf (A) and 6 dpf (B). (C-D) Glomerular *fgfr1a* isoform expression based on RNA-Seq data 5 dpf and 6 dpf. (E) Schematic overview of the *fgfr1a* isoforms found in zebrafish glomeruli. Blue: coding exons; orange: untranslated regions (UTRs). (F) Alignment of the non-equal protein sequences of the two fgfr1a protein isoforms. (G) Protein structural analyses between the two Fgfr1a isoforms 204 (ENSDART00000147742) and 201 (ENSDART00000024882) (red = no change; cyan = insertion; black = substitution; blue = deletion; gray = interaction chain to FGF2). All gene annotations used in this analysis were based on Ensembl release 113 (October 2024).

Furthermore, our results indicate that treatment of zebrafish larvae with NFP disrupts splicing homeostasis. RNA sequencing of both whole larvae and isolated glomeruli revealed an upregulation of serine/arginine-rich splicing factors (SRSFs) like *srsf3a* at 5 and 6 dpf, whereas *srsf2a*, *srsf2b*, and *srsf4* were downregulated (Fig. 7A and Fig. 7F and 7I). Interestingly, most heterogeneous nuclear ribonucleoproteins (hnRNPs; components of the spliceosome) showed consistent downregulation (Fig. 7A).

**Fig. 7:**
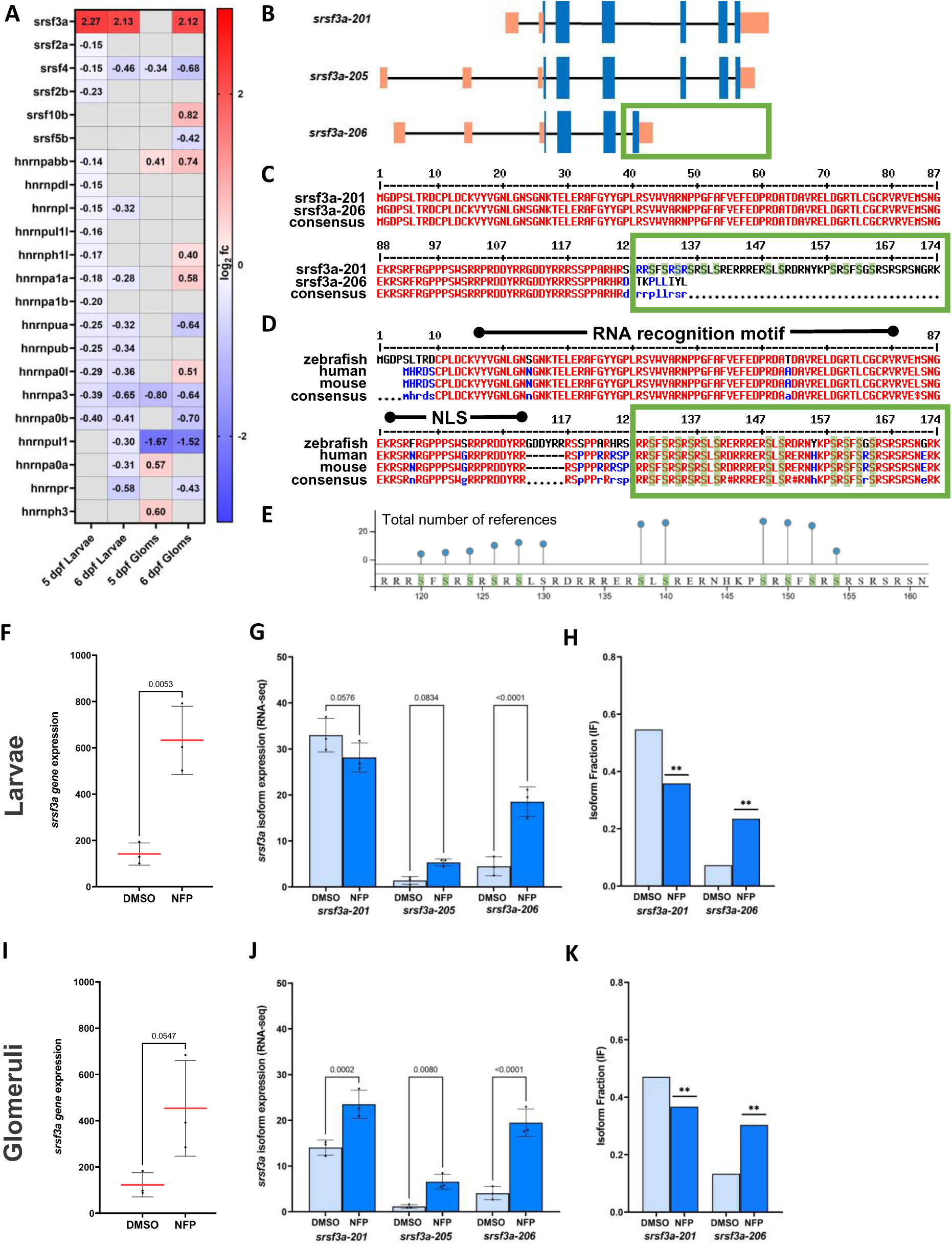
**Correlations between Splicing Factors and differential splicing events.** (A) Heatmaps of gene expression levels of the SRSFs and hnRNPs genes. Data are represented as log2 fold change, a p-value < 0.05 was considered as significant. Red: up-regulated; blue: down-regulated compared to controls. (B) Overview of the *srsf3a* isoforms found in zebrafish larvae and isolated glomeruli. Blue: coding exons; orange: untranslated regions (UTRs). (C) Alignment of the protein sequences of srsf3a-201 and srsf3a-206 isoform. (D) Alignment of the SRSF3 protein sequences between zebrafish, human and mouse. Known phosphorylations sites are marked in green. (E) Published and described phosphorylation sites of Srsf3 from PhosphoSitePlus^76^. (F, I) Gene expression of *srsf3a* in larvae (F) and isolated glomeruli (I) in DMSO and NFP samples at 6 dpf. (G, J) *Srsf3a* isoforms expression in zebrafish larvae (G) and in glomeruli (J) at 6 dpf. Gene counts from RNA-Seq were normalized using DESeq2. (H, K) *Srsf3a* isoform fraction in larvae (H) and glomeruli (K) at 6 dpf based on IsoformSwitchAnalyzeR. All gene annotations used in this analysis were based on Ensembl release 113 (October 2024).

Two *srsf3a* isoforms (Fig. 7B) could be confirmed, with a significant upregulation and higher usage of *srsf3a* isoform 206 (ENSDART00000154963) and a lower usage of isoform srsf3a-201 (ENSDART00000028217) after NFP treatment to induce FSGS (Fig. 7F-K). Both isoforms retain the RNA recognition motif and their nuclear localization signal (NLS) (Fig. 7D), indicating that nuclear translocation and RNA binding are likely not impaired. However, a notable difference is the absence of a large C-terminal region comprising 38 amino acids in isoform 206 (Fig. 7B-C). Interestingly, this C-terminal region – which is highly conserved across zebrafish, humans, and mice – contains a higher density of serine residues (Fig. 7D-E). Twelve serine positions within this region have been repeatedly described in the literature as phosphorylation sites (Fig. 7E). This suggests that activation of *srsf3a* may occur within this domain, a process that is likely disrupted in isoform 206, which increases in the FSGS zebrafish model.

## Discussion

In this study, we performed RNA-Seq analysis to investigate alternative splicing in the pathogenesis of focal segmental glomerulosclerosis (FSGS) using a transgenic zebrafish model. Administration of Nifurpirinol (NFP) successfully induced FSGS in zebrafish larvae, including podocyte depletion, periocular and abdominal edema, and altered glomerular gene expression such as changes in nephrin, podocin, and other glomerular and podocyte markers. Our previous observations have shown that two time points are crucial for studying the progression of FSGS in zebrafish larvae: 5 dpf, representing an early stage of the disease, and 6 dpf, indicating an advanced stage.^11–13^ Accordingly, a time-dependent increase in the number of differentially expressed and differentially spliced genes was observed. This analysis highlights the dynamic nature of transcriptomic changes during disease progression.

Our cross-species analysis underscores the translational validity of the zebrafish FSGS model by revealing substantial overlap in glomerular gene expression changes with human FSGS datasets. Notably, key podocyte markers (e.g. *nphs1*, *nphs2*, *wt1*) and transcription factors (*foxc1*, *mafb*) were downregulated, consistent with previous human and murine FSGS studies.^36–42^ The upregulation of *stat3*, a well-known driver of podocyte injury and glomerulonephritis,^32^ further supports the relevance of our model for studying inflammatory signaling in FSGS. Moreover, elevated expression of extracellular matrix genes such as *fn1* and *serpine1* aligns with known podocyte-associated stress responses.^9,43,44^ The upregulation of pro-inflammatory mediators *il6r* and *il1st* reflects conserved activation of cytokine signaling.^45^ In this context, the observed upregulation of *il6r*, *il6st*, and *jak1* likely contributes to enhanced STAT3 activation through increased phosphorylation, promoting its nuclear translocation and induction of downstream targets such as SOCS3.^46^ Simultaneously, downregulation of *ptpro* may impair STAT3 dephosphorylation, further sustaining its activation. PTPRO functions as a protein tyrosine phosphatase and has been shown to directly dephosphorylate STAT3 at Tyr705, thereby inhibiting its activity.^47,48^ Given the established role of STAT3 activation in kidney injury, particularly in podocytes and tubular epithelial cells, a PTPRO-mediated inhibition of STAT3 could exert protective effects.

Additionally, several microRNAs (miRNAs) known to be involved in glomerular diseases were detected in our zebrafish FSGS model, highlighting their potential relevance to disease pathogenesis. miR-21, which could potentially serve as a biomarker for FSGS and miR-192, which is also associated with podocyte injury, were significantly upregulated after induction of glomerular injury.^33,49,50^

Notably, miRNA-193, which was also significantly upregulated in FSGS zebrafish larvae, plays a crucial role in the pathogenesis of FSGS.^34,51^ In transgenic mouse models, its overexpression induces podocyte injury by downregulating Wilms tumor protein 1 (*Wt1*), leading to cytoskeletal destabilization and glomerular sclerosis.^34^ Its clinical association with primary FSGS and diagnostic potential via urinary and plasma biomarkers underscores its relevance.^52,53^ Targeting miR-193 may offer new therapeutic strategies for FSGS and chronic kidney disease.^54^

Subsequently, we investigated alternative splicing patterns in zebrafish glomeruli affected by FSGS. Our analysis revealed differential exon usage in key podocyte-specific genes, including *nphs1*, *nphs2*, *podxl*, *ptpro*, and *magi2*. Previous studies have provided evidence that these genes are subject to alternative splicing.^55–61^ Nonetheless, the functional implications of these splicing events remain to be elucidated and warrant further investigation.

Exon skipping was the most common type of AS identified in this study. This result is in line with previous studies in humans and animals, confirming this evidence also in zebrafish.^62,63^ A more detailed analysis of alternatively spliced genes in zebrafish glomeruli led to the identification of three potentially important genes: *epb41l5*, *fgfr1a,* and *srsf3a*. *Epb41l5*, highly expressed in podocytes, is critical for focal adhesions; its deletion in models causes proteinuria and FSGS-like damage.^35,64–66^ *Fgfr1*, also enriched in podocytes and PECs, is involved in cell signaling, with dysregulation linked to renal disease.^67–69^ However, the splicing events and isoform switch of these genes have not been described in the field of renal disease, particularly FSGS. While many studies focus on looking at ’global’ gene expression, our study highlights the importance of analyzing individual isoforms, as they may be regulated differently from their global gene expression counterparts. Several isoforms show the phenomenon known as ’isoform switch’ between control and disease samples.^70–72^ Therefore, isoforms can provide a more detailed view of the functional implications in the context of the diseases.

The splicing mechanism of pre-mature RNA is mainly regulated by two families of proteins: hnRNPs (heterogeneous nuclear ribonucleoproteins) and SRSFs (serine-/arginine-rich splicing factors, or SR proteins). We observed dysregulation of both families in our zebrafish FSGS model. SR proteins primarily recruit spliceosome components to promote exon inclusion, whereas hnRNPs inhibit splice site recognition and subsequent exon inclusion. An alteration in the balance between hnRNPs and SRSFs may be associated with disease.^73,74^

Certain limitations of this study should be considered. Although the zebrafish model recapitulates key features of human FSGS, including foot process effacement, podocyte detachment, matrix accumulation, proteinuria and activation of the parietal epithelial cells as well as zebrafish genes share high homology with those of mice and humans, species-specific differences in gene regulation and glomerular architecture cannot be fully excluded. Nevertheless, the high reproductive capacity of zebrafish and their suitability for rapid, large-scale screening make them a valuable system to explore disease mechanisms with potential relevance for higher vertebrates.

In conclusion, our findings highlight the regulation of gene and miRNA expression as well as the importance of AS in the pathogenesis of FSGS suggesting that targeting (splicing) dysregulation could be a promising therapeutic strategy. The combination of transcriptomic analysis and the zebrafish model provides a powerful platform for identifying novel disease mechanisms and potential treatments for kidney diseases such as FSGS.

## Author Contributions Statement

The study was designed by K.E., N.E., U.V., J.B., and F.K.; F.K., F.M. contributed to the zebrafish experiments; RNA-Seq was performed by S.F.; RNA-Seq data were analyzed by O.T. and S.S.; protein structure analysis was done by A.G. and O.K., miRNA experiments were performed and analyzed by A.I. and T.L.; single cell RNA-Seq data were analyzed by F.S.; all other experiments were performed by F.K. and F.M.; experimental data were analyzed by F.K., F.M., E.H., O.T., S.A., S.S.; T.K. consulted on data analysis; F.K., F.M. and N.E. wrote the main manuscript text. F.K. and F.M. prepared figures. All authors reviewed the manuscript.

## Data availability

RNA-Seq data are available in Supplementary Data 1. A comprehensive list of all identified AS genes is provided in Supplementary Data 3. The RNA-Seq used in the study data (the raw data and the preprocessed data) has been deposited at the GEO database with restricted access. Reviewers can access the data using the reviewer token: https://www.ncbi.nlm.nih.gov/geo/query/acc.cgi?acc=GSE301334. All other relevant data and materials are available within the paper and its Supplementary Information and from the corresponding author on reasonable request.

## Acknowledgements

The authors thank Anja Wiechert, Claudia Weber, and Jurij Barmenkov for technical assistance as well as Oliver Zabel and Steffen Prellwitz for the excellent zebrafish husbandry.

This work was supported by grants of the Federal Ministry of Education and Research to Uwe Völker and Nicole Endlich (BMBF, grant 01ZX1908B + 01ZX2208B, Sys_CARE), Prof. Dr. Olga Kalinina (BMBF, grant 01ZX2208C and 01ZX1908C) and to Jan Baumbach (BMBF, grant 01ZX1908A and 01ZX2208A). This work was generously supported by the Südmeyer fund for kidney and vascular research (‘Südmeyer Stiftung für Nieren- und Gefäßforschung’) and the Dr. Gerhard Büchtemann fund, Hamburg, Germany. The funders had no role in study design, data collection and analysis, decision to publish, or preparation of the manuscript.

---

**Fig. S1:**
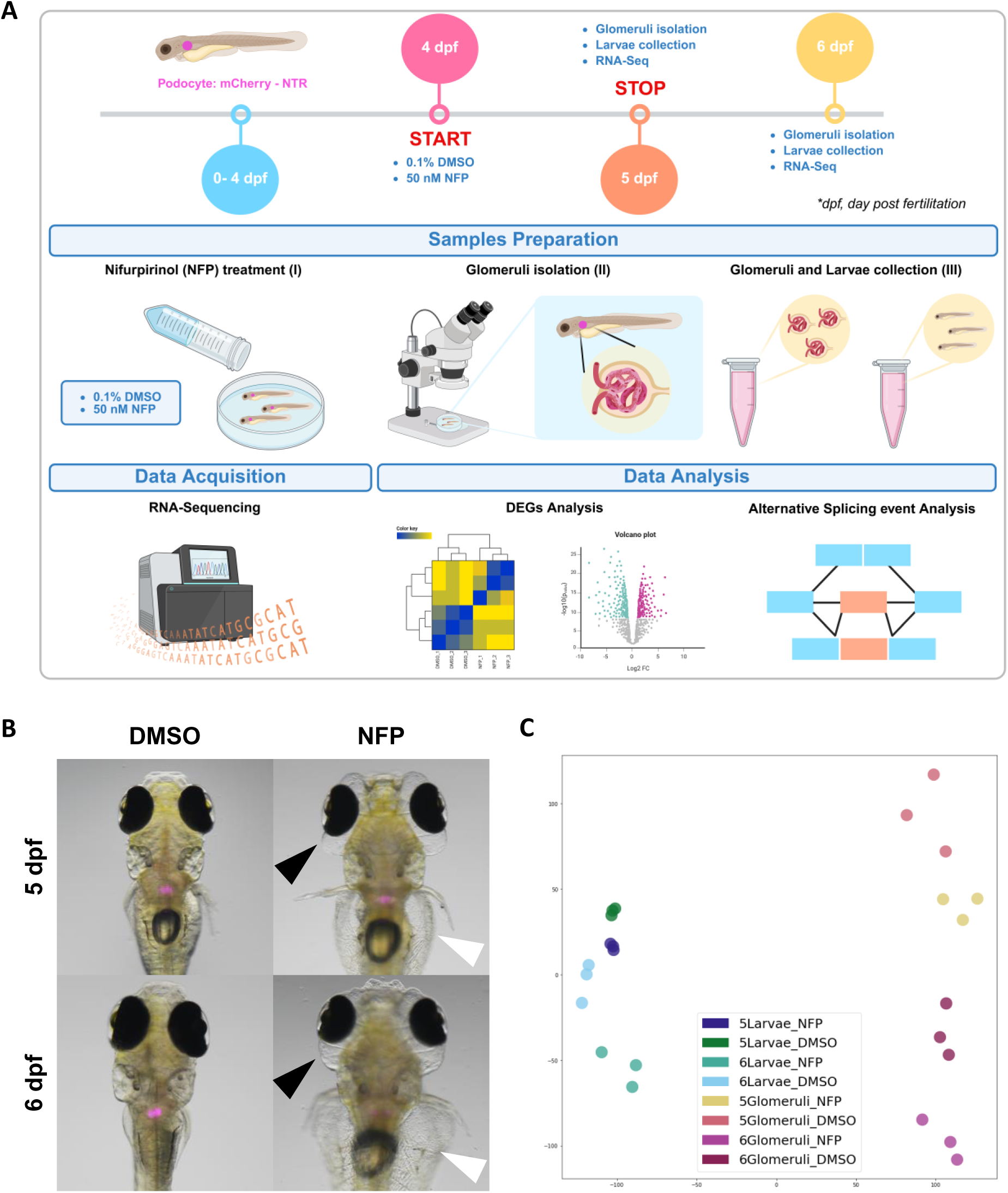
**Overview of the experimental design of the FSGS model in Zebrafish larvae.** (A) Representation of the experimental design with: timeline, sample preparation and data acquisition. (B) Control-treated and NFP-treated zebrafish mCherry larvae at 5 and 6 dpf. The black arrowhead indicates periocular edema, the blank arrowhead indicates abdominal edema. (C) Scatter plot shows the result of Principal Component Analysis (PCA) of gene expression levels of NFP- and DMSO treated larvae and glomeruli at 5 and 6 dpf. Sample classes (as set with the interesting groups argument) are highlighted in different colors.

**Fig. S2:**
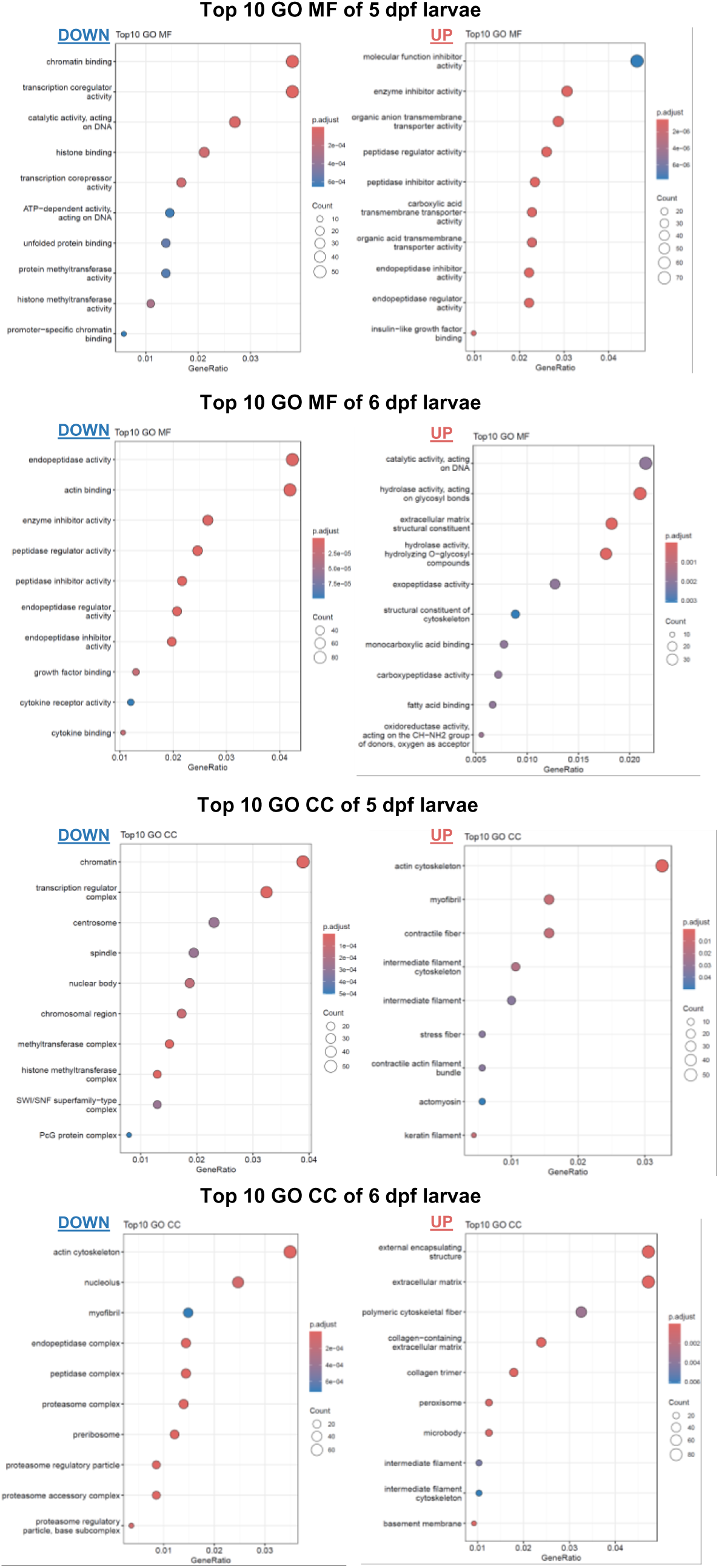
**Detailed gene set enrichment analysis (GSEA) of significantly regulated genes (transcriptome based) in FSGS zebrafish larvae.** RNA-Seq based enriched Top10 GO (Gene Ontology) clusters of significantly regulated genes in FSGS zebrafish larvae. Demonstrated are GO clusters from 5 dpf and 6 dpf larvae categorized in molecular function (MF) cellular components (CC).

**Fig. S3:**
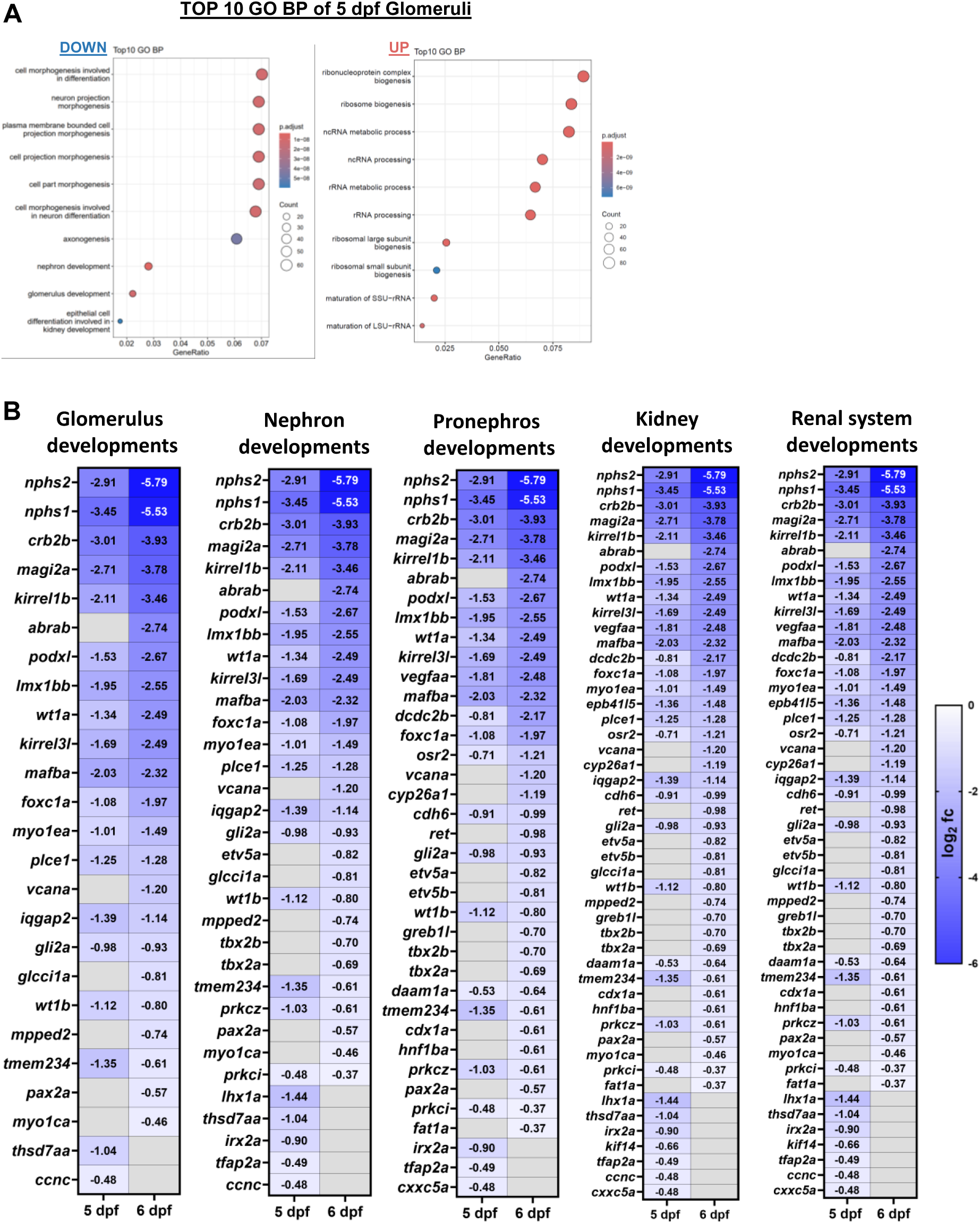
**Detailed gene set enrichment analysis (GSEA) of significantly regulated genes (transcriptome based) in FSGS zebrafish glomeruli, including heatmaps of glomerulus-specific genes.** (A) RNA-Seq based enriched Top10 GO (Gene Ontology) clusters of significantly regulated genes in FSGS zebrafish glomeruli. Shown are GO clusters from 5 dpf zebrafish glomeruli categorized in biological processes (BP). (B) Heatmap showing DEGs in isolated zebrafish glomeruli at 5 and 6 dpf, identified by DESeq2. Gene clusters related to glomerulus, nephron, pronephron, kidney and kidney system developments are shown.

**Fig. S4:**
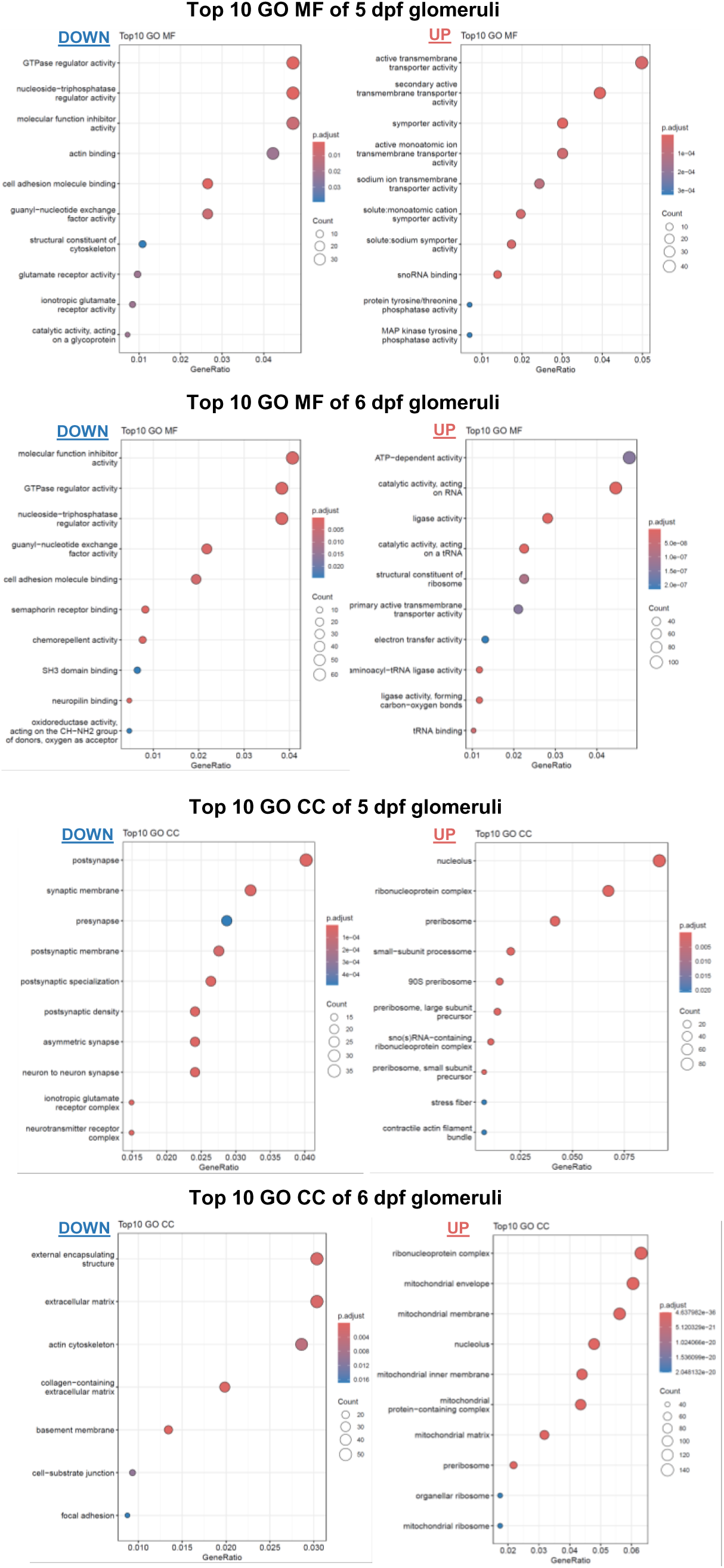
**Detailed gene set enrichment analysis (GSEA) of significantly regulated genes (transcriptome based) in FSGS zebrafish glomeruli.** RNA-Seq based enriched Top10 GO (Gene Ontology) clusters of significantly regulated genes in FSGS zebrafish glomeruli. Demonstrated are GO clusters from 5 dpf and 6 dpf zebrafish glomeruli categorized in molecular function (MF) cellular components (CC).

**Fig. S5:**
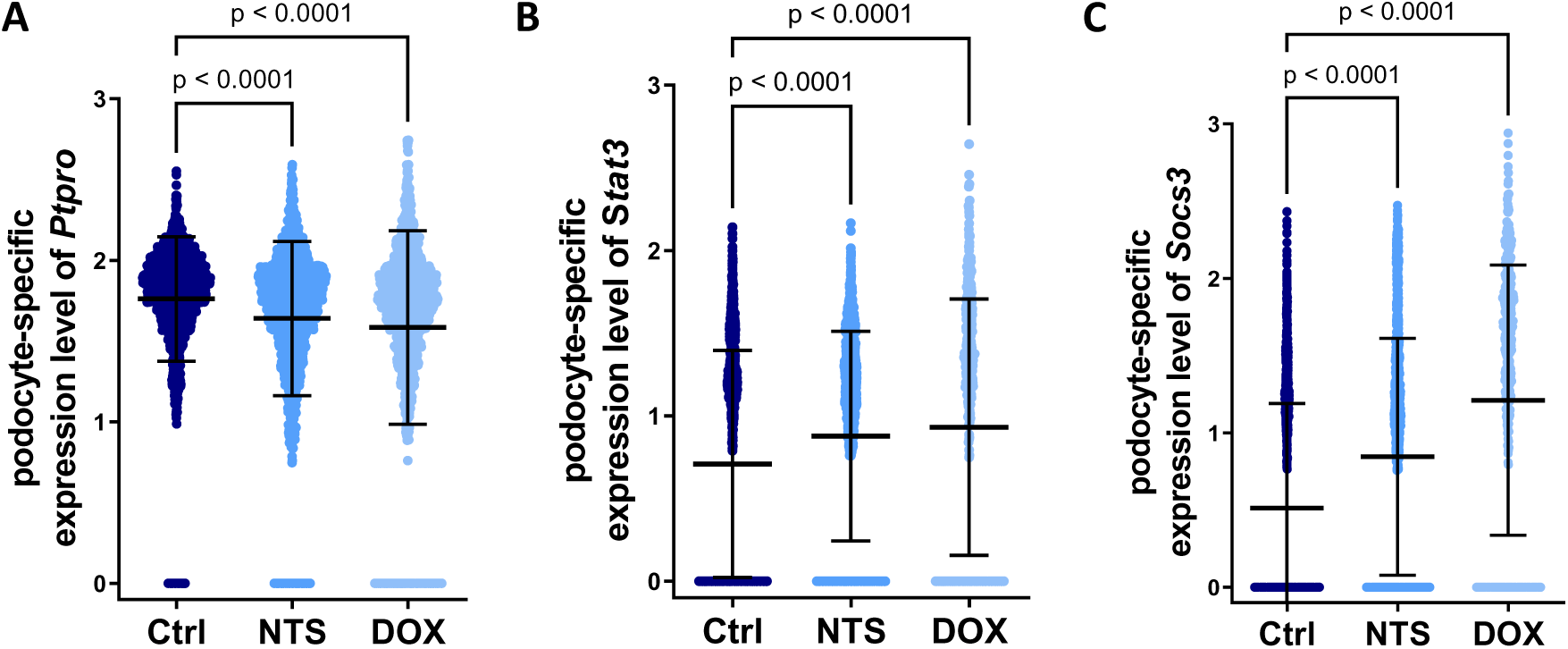
Single-cell RNA-Seq analysis of *Ptpro*, *Stat3* and *Socs3* expression in different kidney injury models. Single-cell RNA-seq data from (Chung et al.^30^), reveal a significant altered podocyte-specific gene expression of (A) *Ptpro*, (B) *Stat3* and (C) *Socs3* in mouse glomeruli following nephrotoxic serum nephritis (NTS) or doxorubicin (DOX) treatment compared to the control (Ctrl).

**Fig. S6:**
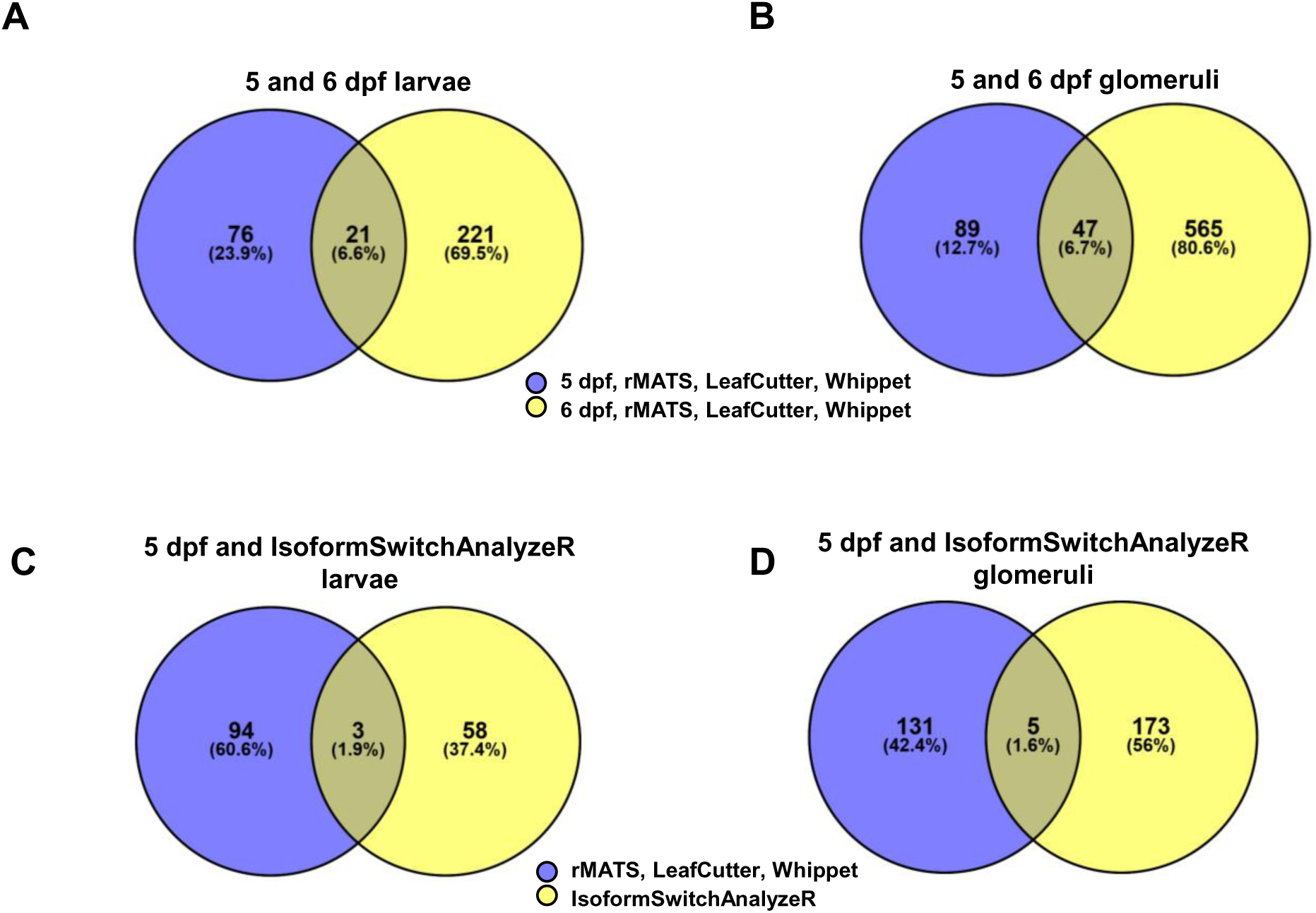
**Detection of alternative splicing events using multiple analysis tools.** Venn diagrams showing alternatively spliced genes using rMATS, LeafCutter, Whippet and IsoformSwitchAnalyzeR in larvae (A, C) and isolated glomeruli (B, D) at 5 and 6 dpf. A comprehensive list of all identified alternatively spliced genes is provided in Supplementary Table 3.

**Fig. S7:**
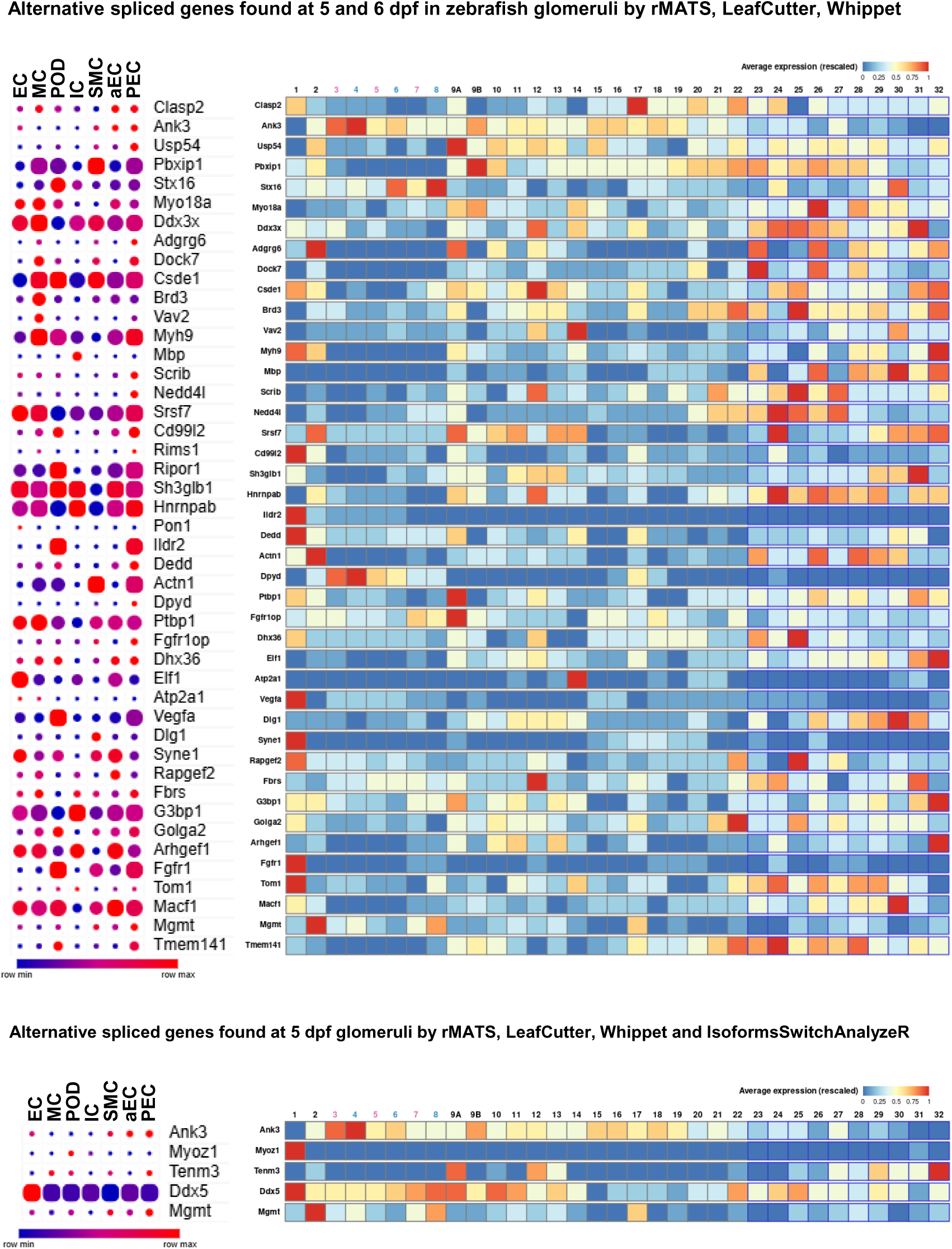
Kidney expression pattern of RNA-Seq based alternative spliced candidates. Single-cell RNA-Seq data of murine kidneys^77^ and mouse glomerulus^30^ to determinate podocyte-specific expression of alternatively spliced genes of Supplemental Figure S6B and S6D. Color code: Red means a high average expression; Blue: low expression. Column caption: (1) podocytes (visceral epithelium), (2) parietal epithelium, (3) segment one of proximal tubule - female, (4) segment one of proximal tubule - male, (5) segment two of proximal tubule - female, (6) segment two of proximal tubule - male, (7) segment three of proximal tubule - female, (8) segment three of proximal tubule - male, (9A) LOH thin descending limb of inner stripe of outer medulla of cortical nephron, (9B) LOH thin descending limb of inner stripe of outer medulla of juxtamedullary nephron, (10) upper LOH thin descending limb of inner medulla of juxtamedullary nephron, (11) lower LOH thin descending limb of inner medulla of juxtamedullary nephron, (12) lower LOH thin limb of inner medulla of juxtamedullary nephron, (13) lower LOH thin limb of inner medulla of juxtamedullary nephron, (14) upper LOH thin ascending limb of inner medulla of juxtamedullary nephron, (15) distal straight tubule of inner stripe of outer medulla (syn: thick ascending limb of LOH), (16) thick ascending limb of LOH), (17) macula densa, (18) distal convoluted tubule, (19) nephron connecting tubule, (20) principal-like cell of nephron connecting tubule, (21) intercalated type non-A non-B cell of nephron connecting tubule, (22) intercalated type A cell of nephron connecting tubule and cortical collecting duct, (23) principal-like cell of cortical collecting duct, (24) intercalated type B cell of cortical collecting duct, (25) intercalated type A cell of outer medullary collecting duct, (26) principal cell of outer medullary collecting duct, (27) intercalated type A cell of inner medullary collecting duct, (28) principal cell of inner medullary collecting duct type one, (29) principal cell of inner medullary collecting duct type two, (30) principal-like cell of deep inner medullary collecting duct type one, (31) cell of deep inner medullary collecting duct type two, (32) deep medullary epithelium of pelvis. EC, glomerular endothelial cells; MC, mesangial cells; POD, podocytes; IC, immune cells; SMC, smooth muscle cells; aEC, arteriolar endothelial cells; PEC, parietal epithelial cells.

**Fig. S8:**
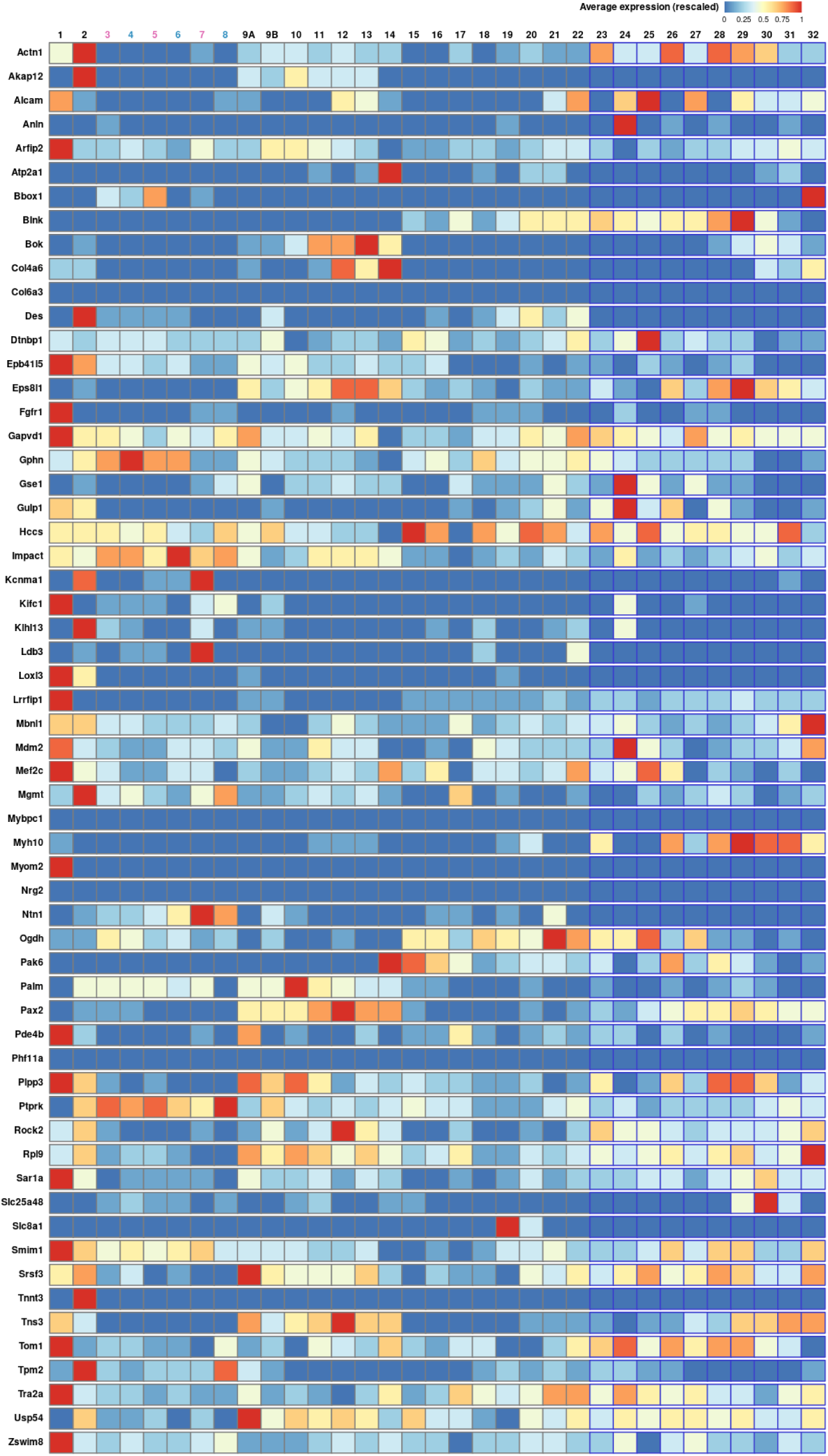
Kidney expression pattern of RNA-Seq based alternative spliced candidates. Database analysis using the Kidney Cell Explorer by Ransick et al.^77^ based on a single-cell RNA sequencing data set of murine kidneys showed the expression pattern for all kidney cell fractions of alternatively spliced genes of figure 4J. Color code: Red means a high average expression; Blue: low expression. Columns are labeled as in Figure S7.

**Fig. S9:**
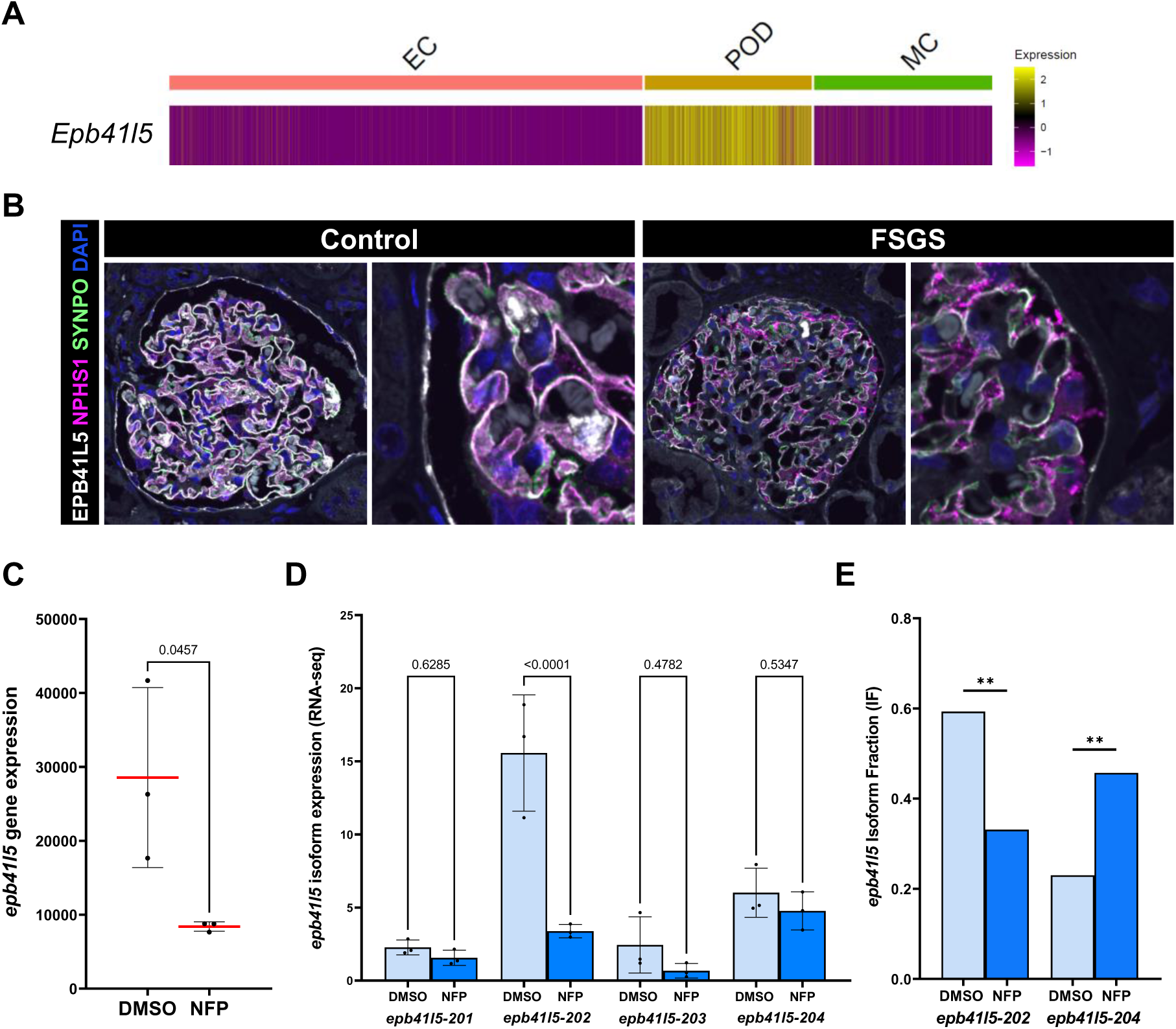
Glomerular *Epb41l5* expression in mice, human and zebrafish. (A) Heatmap of single-cell RNA expression data of *Epb41l5* in mice glomeruli showed significant enrichment in the podocyte cluster (POD). Color code: Yellow means a high average expression; magenta: low expression. EC = endothelial cells; MC = mesangial cells. (B) Control and FSGS human kidney sections were stained with antibodies against EPB41L5 (shown in white), nephrin (magenta) and synaptopodin (green). Nuclei were stained with DAPI (blue). (C) Gene expression of *epb41l5* in zebrafish glomeruli treated with NFP for FSGS induction at 5 dpf confirmed *epb41l5* downregulation after FSGS. (D) Expression of *epb41l5* isoforms based on DESeq2 normalized gene counts from RNA-Seq at 5 dpf. (E) *Epb41l5* isoform fraction based on IsoformSwitchAnalyzer data at 5 dpf.

